# Soil bacterial community diversity and composition vary more with elevation than seasons in alpine habitats of western Himalaya

**DOI:** 10.1101/2020.11.05.370114

**Authors:** Pamela Bhattacharya, Samrat Mondol, Gautam Talukdar, Gopal Singh Rawat

## Abstract

Soil heterotrophic respiration-driven CO_2_ emissions, its impact on global warming and the mechanistic roles of soil bacterial communities in this process have been an area of active research. However, our knowledge regarding the effects of environmental changes on soil bacterial communities is limited. To this end, the climate-sensitive high-altitude alpine ecosystems offer ideal opportunities to investigate relationship between climate change and bacterial communities. While data from several high-altitude mountain regions suggest that local environment factors and geological patterns govern bacterial communities, no information is available from the Himalaya. Here we provide baseline information on seasonal soil bacterial community diversity and composition along a 3200-4000 m elevation gradient covering four alpine habitats (subalpine forest, alpine scrub, alpine meadow and moraine) in Gangotri National Park, western Himalaya. Bacterial metabarcoding data from 36 field-collected samples showed no elevation trend in the bacterial richness and a non-monotonous decrease in their diversity. Further, their community diversity and composition varied significantly among habitats along elevation but were stable seasonally within each habitat. The richness was primarily influenced by soil inorganic carbon (SOC) and total nitrogen (TN), whereas temperature, SOC and TN affected diversity and composition patterns. Given the importance of the Himalaya in the context of global carbon cycle this information will help in accurate modeling of climate adaptation scenarios of bacterial niches and their downstream impacts towards climate warming.

## Introduction

Climate warming is one of the major global challenges significantly impacting species, society and ecosystems [1]. Increasing natural and anthropogenic emissions of greenhouse gases, particularly carbon dioxide (CO_2_) play a major role in global warming [2, 3]. Among the natural sources of CO_2_ emission, soil respiration (both autotrophic and heterotrophic) accounts for the largest land to air carbon flux (annually ~90 Pg, nine times higher than anthropogenic efflux) [2, 4–6]. However, this loss of carbon is superseded by photosynthetic inputs leading to overall increase in soil carbon content (>2000 Pg globally) [7, 8]. Any shift in this balance due to climatic factors (temperature, precipitation, increasing CO_2_ levels, N-deposition etc.) may result in a positive feedback to climate warming [6, 9]. In this context, the soil bacterial communities play a key role in CO_2_ emission through heterotrophic respiration [5, 10]. Despite their known mechanistic role in CO_2_ emission and susceptibility to climatic conditions [11], knowledge regarding response of soil bacterial communities to environmental changes is limited [12]. Recent carbon dynamics-based ecosystem models with bacterial community compositions from various trophic levels have shown better predictive power for global soil carbon cycle [7, 13–15]. This underscores the need for studies on soil bacterial community response to environmental change especially in the climate sensitive ecosystems in order to understand and predict alterations in carbon flux globally.

Alpine ecosystems in the high mountains are considered among the most climate sensitive regions [16–18]. These ecosystems are characterized by large carbon sinks (high soil organic carbon (SOC), [19], low decomposition and turn over rates [19] but high bacterial diversity [20]. They also exhibit strong seasonal and annual fluctuations in environmental conditions along elevation gradient [21] across various habitats. These features make the alpine habitats as ideal systems to investigate the relationship between the prevailing climatic conditions and bacterial community [16, 20]. Some of the recent studies have generated data on bacterial community composition along elevation gradient in different alpine habitats of the world, analyzed impacts of seasonal variation on them and found area-specific patterns (see Supplementary Table S1). For example, a number of alpine habitats from Swiss and Italian Alps have shown decreasing [20, 22] as well as no specific bacterial richness and diversity trends along elevation [23, 24]. In contrast, sites from Mount Wutai and Tibetan Plateau, China have shown increasing trend in bacterial richness and diversity [25, 26], whereas sites in Changbai Mountain and Mount Gongga reported a decreasing trend (Supplementary Table S1). These studies also reported habitat-specific bacterial community compositions [20, 22] and observed seasonal variations in them (Supplementary Table S1). All of this information suggests that soil bacterial communities are influenced by local environmental factors (for example mean annual temperature, mean annual precipitation, soil pH, soil organic carbon and nitrogen, C/N ratio etc.) governed by geology and thus need to be investigated at local/regional scales. While such information is available from various high altitude mountain regions of the world (see Supplementary Table S1), data from the Himalayan region is lacking [27, 28].

In this paper, we address this gap in our knowledge by reporting soil bacterial communities and their relationship with various environmental factors along an elevation gradient and across seasons (spring and autumn) covering four habitats in the alpine region of western Himalaya, India. The Himalaya, representing diverse ecosystems is the highest mountain range in the subcontinent comprising diverse ecosystems that are extremely sensitive to climate change [29–31]. It retains 33% of India’s carbon stock [32] and is known to play a critical role in global carbon cycle [30]. Recent assessments suggest regular fluctuations in precipitation and project an increase of 3°C temperature by 2050 across Himalaya [33], resulting in destabilization of soil organic matters [31]. We collected environmental and bacterial community data along an elevation from Gangotri National Park (GNP), western Himalaya. This park is represented by varying topography and diverse habitats [34]. Main objectives of the study were (i) characterization of environmental variables across four habitats viz. subalpine forest, alpine scrub, alpine meadow and moraine along elevation gradient and seasons (Spring and Autumn); (ii) assessment of bacterial community richness, diversity and composition; and (iii) evaluation of the relationship between the environmental variables and bacterial community diversity and composition. The results provide the first baseline information on the Himalayan alpine soil bacterial community and their relationship with local environmental variables.

## Materials and Methods

### Research permissions and ethical considerations

Permissions for field surveys and soil sampling were provided by Uttarakhand Forest Department (Permit no. 702/5-6). Soil sampling did not require any specific ethical clearance.

### Study Area

We conducted this study in Gangotri National Park (GNP), the largest high altitude national park (~2390 sq. km) in Uttarakhand, India (Figure 1). This park is situated between 79° 49’ to 79° 25’ E and 30° 43’ to 31° 28’ N covering two Indian biogeographic zones: Trans-Himalaya and Western Himalaya [34, 35]. The park was established in 1989 to conserve high altitude ecosystems. The topography is of montane type representing high mountain ranges, snow-clad peaks, sharp valleys with heterogeneous slopes, alpine meadows and glacial fore-fields covering an altitudinal range of 2600–7138 m. The climate in this area is characterized by long winters with heavy snowfall, cold summer and short cool spring and autumn. The park receives 90% of the precipitation in the form of rainfall during June to September and heavy snowfall from late December to April [34].

**Figure 1:**
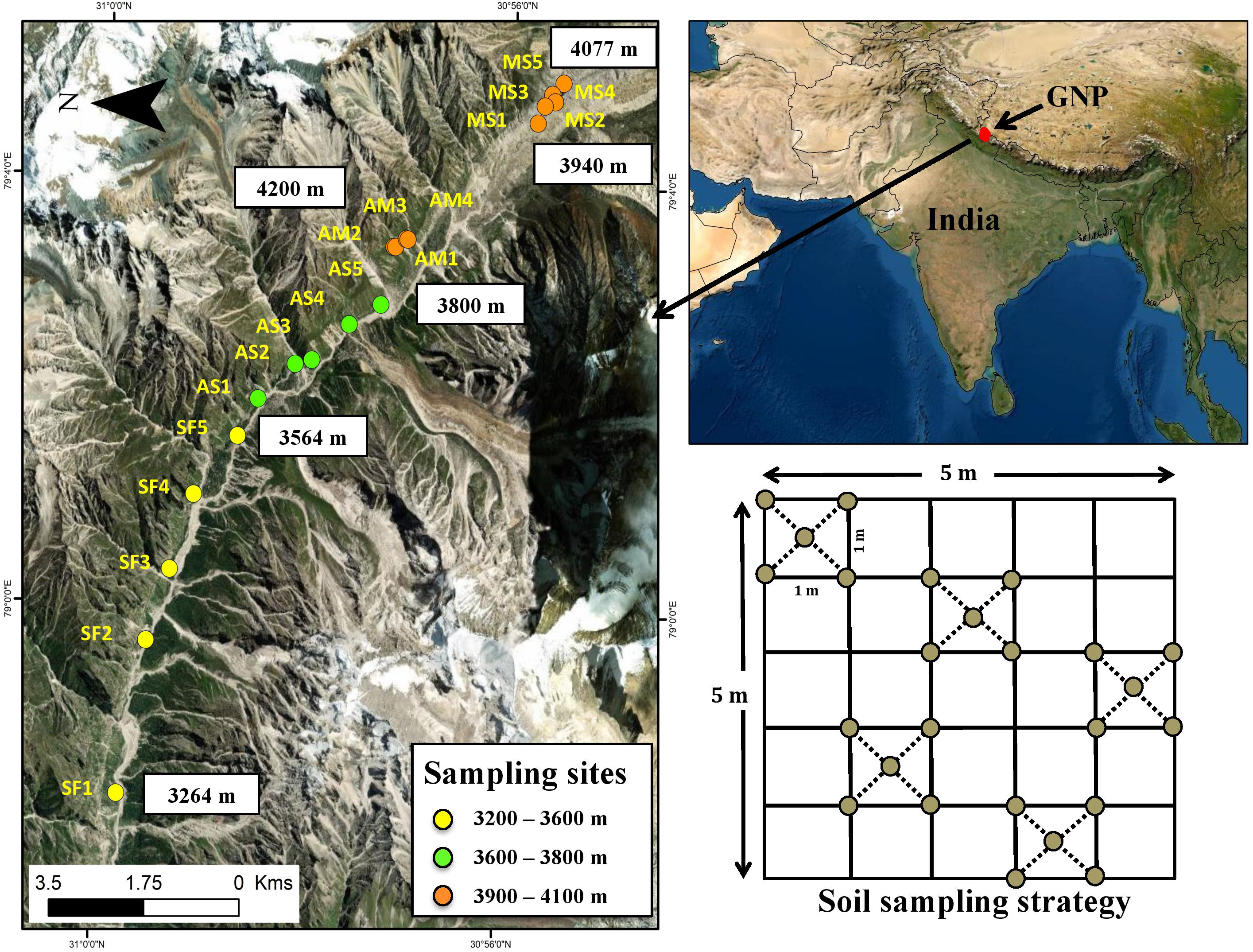
Study area map showing soil-sampling sites covering four habitats along elevation gradient in Gangotri National Park. Site abbreviations: SF-Subalpine forest; AS-Alpine scrub; AM-Alpine meadow; MS-Morainic slope.

Systematic soil sampling was conducted in the upper catchment of Bhagirathi river between Gangotri and Gaumukh i.e., snout of Gangotri glacier (Figure 1) where four distinct habitats: subalpine forest (SF), alpine scrub (AS), alpine meadow (AM) and morainic slope (MS), were clearly discernible within ~900 m elevation gradient (between 3200-4100 m). During the study period (2016-2017), we measured the mean annual air temperature and relative humidity, and found a pattern of decreasing temperature (7.81±0.28°C-1.28±0.12°C) and increasing relative humidity (57.7±3.54%-67.34±3.29%) with increase in elevation (Table 1). We selected 19 sampling sites along a 16 km trail at different elevations located on southwest slope of the valley (Figure 1, Table 1). These sites were divided among four habitats (five sites each in the SF, AS, and MS, whereas four sites in the AM) as independent replicates representing within site variability. AM and MS were located at similar elevation range (3940–4080 m), separated by ~3.5 km distance (Figure 1). The detailed habitat characteristics of the study sites are provided in Table 1.

**Table 1:**
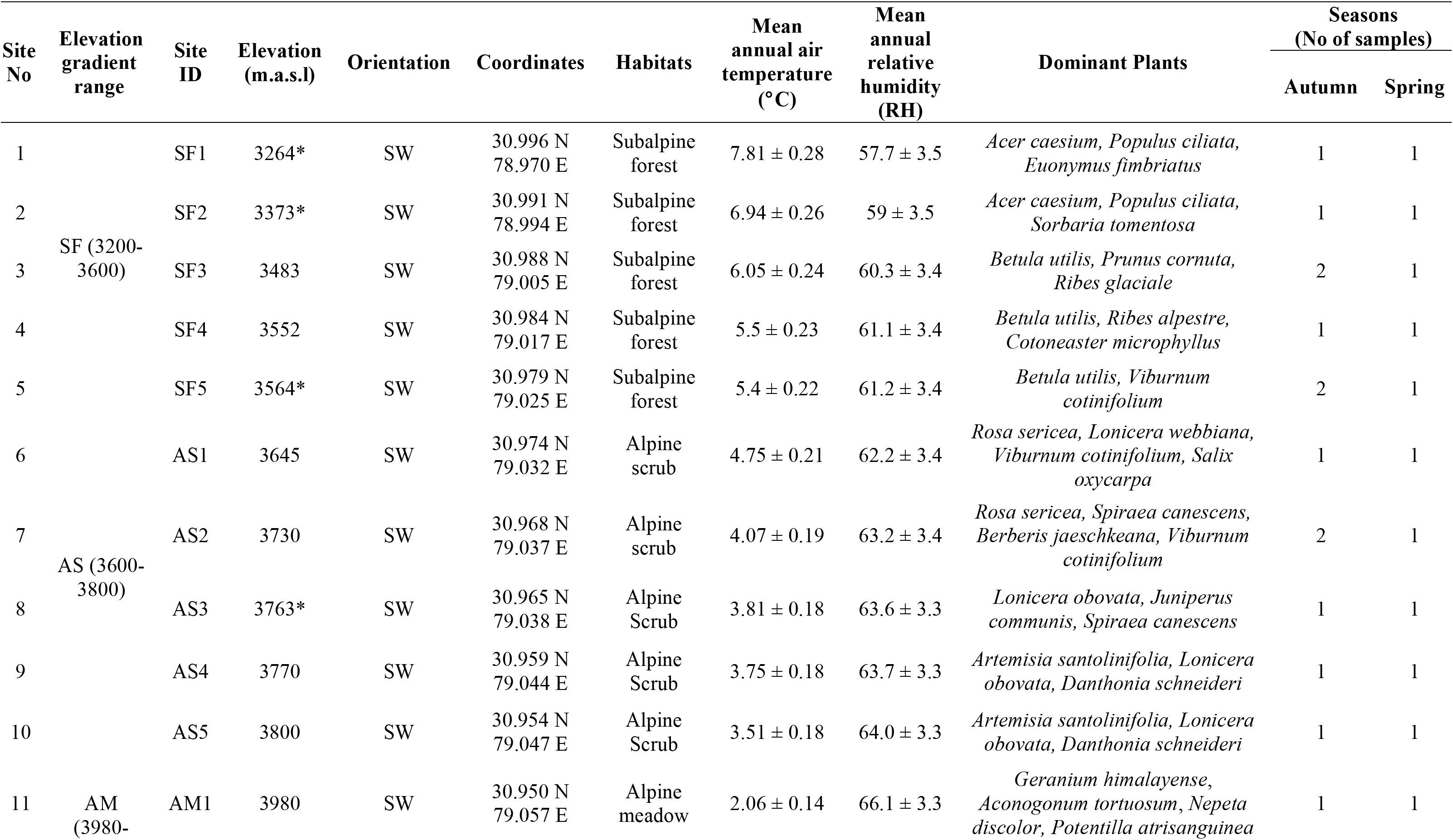
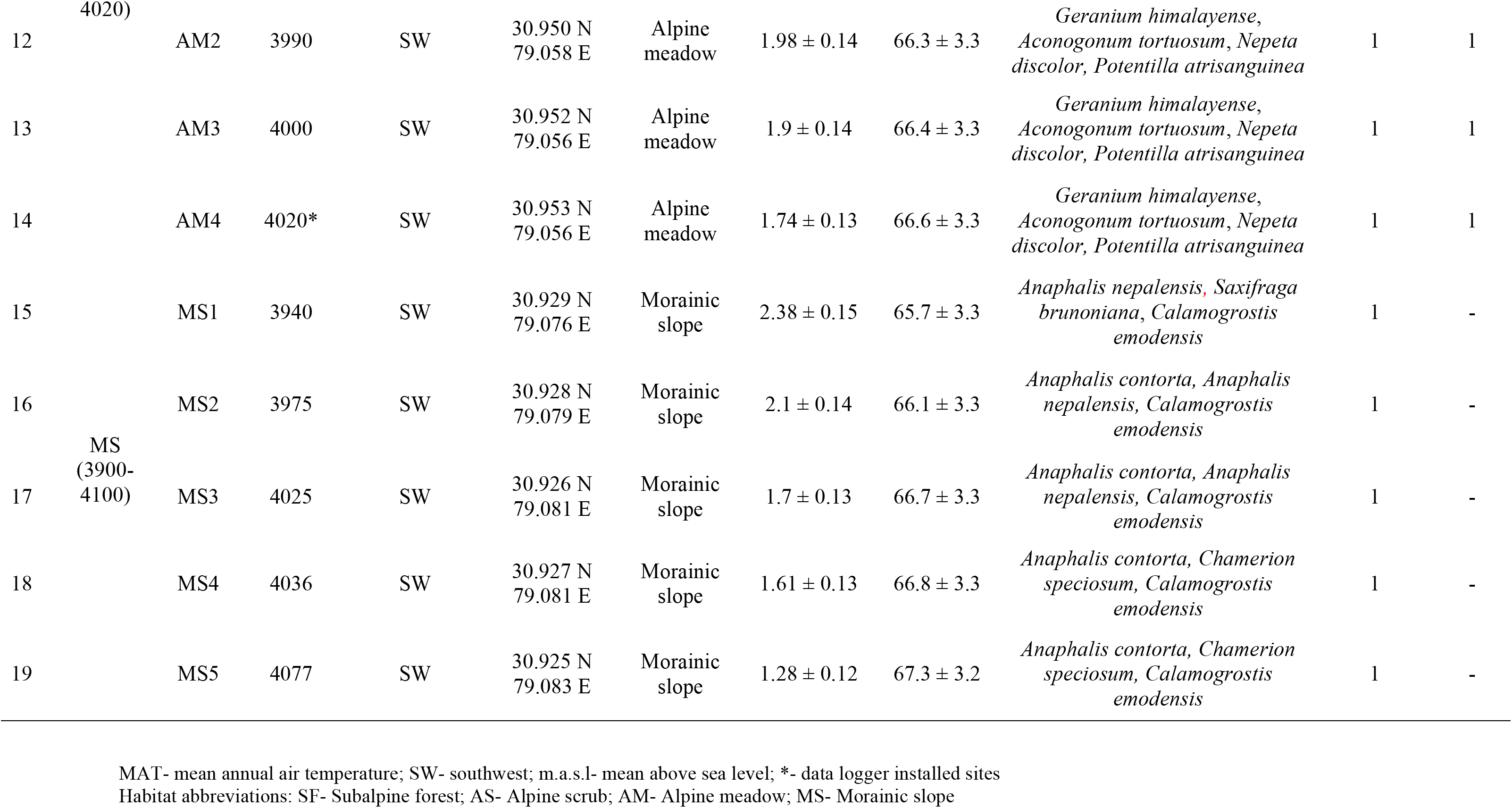
Details of the sampling sites.

### Soil sample collection

We collected soil samples at the selected habitats and seasons (Autumn - October 2016 and early Spring - May 2017) [34]. Spatial sampling was conducted once from all 19 sites during autumn. We collected a total of 22 samples from these four habitats (one sample each from 19 sites along with spatial replicates at three sites). Seasonal sampling was performed within three habitats (SF, AS and AM) during spring, where we collected one sample from the same locations across all sites (n=14). Overall, we collected 36 soil samples for the intensive study area (Table 1).

At each sampling site, we randomly selected five 1 m X 1 m plots within a 5 m X 5 m sampling area as spatial replicates (Figure 1). Plant litters was removed from the surface and five soil cores of approximately 2 g weight/core (four corners and the center of each plot) were collected from top 5 cm using hand held sterile soil corer (50 ml centrifuge tube, Tarsons Products Pvt. Ltd, India) [25]. After collecting soil from four more plots, all samples were pooled (~50 g) in a sterile zip-lock bag and homogenized manually. The homogenized sample pool was subsampled in two aliquots: a ~5 g aliquot, stored in 100% ethanol (Merck, Germany) for molecular analyses and the remaining sample (~45 g) was kept in the zip-lock bag. Both aliquots were stored in a cool box in the field and transported to the laboratory, where the smaller subsamples were stored in a −20°C freezer and the other aliquots were kept at 4°C until further analyses.

### Collection of environmental data

We installed five data loggers (HOBO U23 pro v2, Onset Computer Corporation, USA) in three habitats at different elevations: 3264 m, 3373 m, 3564 m (SF), 3763 m (AS) and 4020 m (AM) to collect air temperature data. Air temperature data were recorded at every one-hour interval during the entire study period. In addition, we also installed two data loggers to measure soil temperature at 3564 m (SF) and 4020 m (AM), respectively. Our analyses show very strong association between daily mean air temperature and soil temperature (Supplementary Figure S1) at 5 cm depth, indicating air temperature data can be used to explain soil bacterial patterns [36]. We used the mean temperature for previous 30-day period (MT_30_) from the soil sampling date for analyses. Based on the temperatures measured at the five data logger points we calculated MT_30_ for all the other soil sampling sites.

We also estimated a number of soil environmental variables such as soil pH, moisture content, organic carbon, total nitrogen and C:N from the field-collected soil samples (n=36). Soil moisture content (SMC) (expressed as percentage of water weight in soils) was measured gravimetrically immediately after transport to laboratory, where 5 g of each sample was weighed before and after drying in oven at 105° C for 48 h [37]. Remaining ~40 g soil was airdried at room temperature and sieved through a 1 mm sieve to remove any remaining large particles, plant roots and leaf litter. Soil pH was measured using a glass electrode pH meter (Sension 7, Hach Company, USA) in a 1:2.5 (w/v) suspension of soil to deionized water that has been vigorously stirred and left to stand overnight [38]. Soil organic carbon (SOC) was measured using the potassium dichromate (K_2_Cr_2_O_7_) oxidation method [39]. The total nitrogen (TN) content was determined by automated Kjeldhal method [40] using Kjeltec 8400 (Foss India Pvt. Ltd, The C:N ratio was calculated by dividing SOC content by total N content.

### DNA extraction, PCR amplification and Illumina sequencing

We extracted soil genomic DNA from approximately 0.25 g of each field-collected soil sample using Nucleopore GDNA soil kit (Genetix Biotech Asia Pvt. Ltd, India) following the manufacturer’s kit protocol. For every set of extraction (n=11 samples) we included one negative control to monitor any possible contamination. To check the DNA quality, we amplified the V3 and V4 region of the bacterial 16S rRNA gene using the primer sequences 338F (5’-CCTACGGGNGGCWGCAG-3’) and 806R (5’-GACTACHVGGGTATCTAATCC-3’) [25]. The PCR reaction contained 5 μl of 2X Kappa Hifi Hotstart ReadyMix (Kappa Biosystems, Roche Sequencing and Life Science, USA), 2 μl of 0.2 mg/ml BSA, 1 μl each of 2.5 μM forward and reverse primer and 1 μl of extracted DNA. PCR conditions included an initial denaturation of 95°C for 3 min, followed by 35 cycles of 95°C for 30 s, 55°C for 30 s, and 72°C for 30 s, followed by a final extension for 10 min at 72°C. PCR negative was included to monitor any contamination. Samples that showed positive amplication (~460bp band) were selected for Illumina sequencing and stored at −20°C until further analysis.

We quantified DNA concentrations of these samples using Qubit dsDNA HS Assay Kit (Life Technologies, USA) and used Illumina 16S Metagenomics Sequencing library preparation protocol to prepare sequencing libraries for V3 and V4 regions. The PCR reaction contained 12.5 μl of 2X Kappa Hifi Hotstart ReadyMix (Kappa Biosystems, Roche Sequencing and Life Science, USA), 1 μl each of 5 μM forward and reverse primer, 10.5 μl of microbial DNA (12.5 ng), respectively. The PCR conditions were same, except for 25 cycles of amplification. The resulting PCR products were run on an Agilent 2100 Bio-analyzer (Agilent Technologies INC., United States) to verify the expected size (~550bp) and were purified using 0.9X Agencourt AMPure XP beads (Beckman Coulter Life Sciences, United States). The second amplification step was carried out using Nextera XT kit (Illumina Inc., United States) with barcode primers, where index PCRs were carried out in 50 μl reactions containing 5 μl of purified DNA, 5 μl each of Nextera XT Index Primer 1 and 2, 25 μl of 2x KAPA Hifi HotStart Ready Mix (Kappa Biosystems, Roche Sequencing and Life Science, USA) and 10 μl of nuclease-free water. PCR conditions were same, except for 8 amplification cycles. Subsequently, amplicons were purified using 1 x Agencourt AMPure XP beads (Beckman Coulter Life Sciences, United States), eluted in 20 μl of 10 mM Tris (pH 8.5), pooled and checked for their size (~ 630 bp) to confirm barcode incorporation. Concentration of amplified products were estimated and normalized (2nM) for each library. Equal volume of these libraries were pooled, denatured and diluted to 4 pM before loading onto the MiSeq flow cell (Illumina Inc., United States). According to Illumina protocol, 12% of PhiX control library was spiked with the amplicon library. Sequencing was performed on Illumina MiSeq platform using a 2×300 bp paired end protocol at the Next Generation Genomics Facility at Center for Cellular and Molecular Platforms (C-CAMP), Bangalore, India.

### Processing of sequence data

We processed the raw sequences using MiSeq SOP pipeline in Mothur v.1.40.5 [41–43]. Fulllength sequences were prepared and good reads (>460 bp length with no homopolymer stretches longer than 8bp) were identified and clustered (based on ≥97% similarity) using 16S rRNA database SILVA 132 SSU SEED [44]. Uchime [45] was used to remove chimeric sequences, followed by removal of all singletons, chloroplasts, archaea, mitochondria and unknown origin sequences. Only high quality sequences were used to estimate pairwise distances and generate single linkage clusters with ≥97% sequence similarity, and the longest read from each cluster was used as the reference sequence for taxonomic assignment against the SILVA SSU NR Ref database [44]. All sequences in each cluster and their replicates (≥97% similarity) provided the quantitative estimates of individual reads per taxonomic unit. The dataset was normalized by rarefying the sequences to lowest sample specific sequencing depth with maximun Good’s coverage [41]. Any sample with sequence read less than the rarefaction depth was discarded. Finally, we clustered the sequences to phylum, class and genera levels for operational taxonomic unit (OTU) abundance, diversity and composition analysis. We further classified the data into abundant and rare taxa based on their relative abundances [46]. OTUs with ≥1% relative abundance were considered abundant, whereas ≤0.1% were considered as rare [47–49].

### Calculations and data analysis

We calculated three bacterial diversity indices: predicted species richness (Chao1), observed species richness (Sobs) and Shannon index (H’) to assess alpha diversity for each sample. Normal distribution and homogeneity of variance of these indices and environmental variables were assessed using Shapiro-Wilk test and Levene’s test, respectively. Since most of the data did not show normal distribution and equal variances, we used Welch’s ANOVA [50] and Mann-Whitney U test to assess variation among habitats and seasons respectively. Further, effect sizes (Cohen’s *d* with hedges correction) were collected to quantify differences in environmental variables and alpha diversity indices [51, 52].

For beta diversity estimation we calculated Bray-Curtis dissimilarity matrix after Hellinger transformation of rarefied abundance data at genus level. To test any significance in compositional differences we used Multi-Response Permutation Procedure (MRPP) analysis [53] with 9,999 permutations, where chance corrected within-group agreement (a measure of effect size presented as *A* value) was calculated. The *A* value ranges between 0 to 1, where *A*=0 means samples within a group are heterogeneous and *A*=1 depicts they are identical. For visual interpretation of site-wise differences in bacterial community composition principal coordinates analysis (PCoA) using Bray-Curtis dissimilarity matrix was used for all, abundant and rare genera, respectively. We performed Similarity Percentage (SIMPER) analysis [54] to identify bacterial phyla, class and genus responsible for the dissimilarity between the bacterial community compositions.

Pearson correlation was used to test the relationship of bacterial richness and alpha diversity with environmental variables. Multiple ordinary least square (OLS) regression analysis was done to assess relationship of environmental variables with bacterial richness and diversity. One of the two highly correlated environmental variables was used in the regression analysis based on their ecological relation. For SOC and TN having high correlation (Supplementary Figure S2), regressions were performed for each separately because of their direct influence on bacterial growth. Mantel test with Pearson correlation was used to test the relationship of bacterial community composition with environmental variables. We performed forward selection of environmental variables using stepwise regression (with 999 permutations) and conducted canonical redundancy analysis (RDA) using Hellinger transformed bacterial abundance data to identify environmental variables that significantly explained the variation in abundant and rare phyla and class. The RDA model and the environmental variables were tested by permutation test using one-way analysis of variance (ANOVA). Finally we used Univariate regression analyses to assess the relationship of environmental variables with Hellinger transformed relative abundance of individual bacterial phyla and class. All analyses were conducted using SPSS software (version 22, IBM, Chicago, IL, USA) and R version 3.2.1 through RStudio (RStudio version 1.2.5042, 2020)[55] with packages, vegan [53], dplyr [56] and ggplot2 [57].

## Results

### Habitat-wise variation in environmental parameters along elevation

During our sampling period, all environmental variables differed along the elevation among the four habitats (Figure 2, Table 2). While mean temperature of 30 days period before sampling (henceforth MT_30_) and C:N ratio showed a clear decreasing trend along the elevation, SOC and TN exhibited overall decreasing trend except high values at AM (Mean_SOC_=60.6 g kg^-1^, SD=20.7, n=8 and MeanτN=5.67 g kg^-1^, SD=1.67, n=8) at higher elevation. The SMC and pH values showed no specific elevation trend.

**Figure 2:**
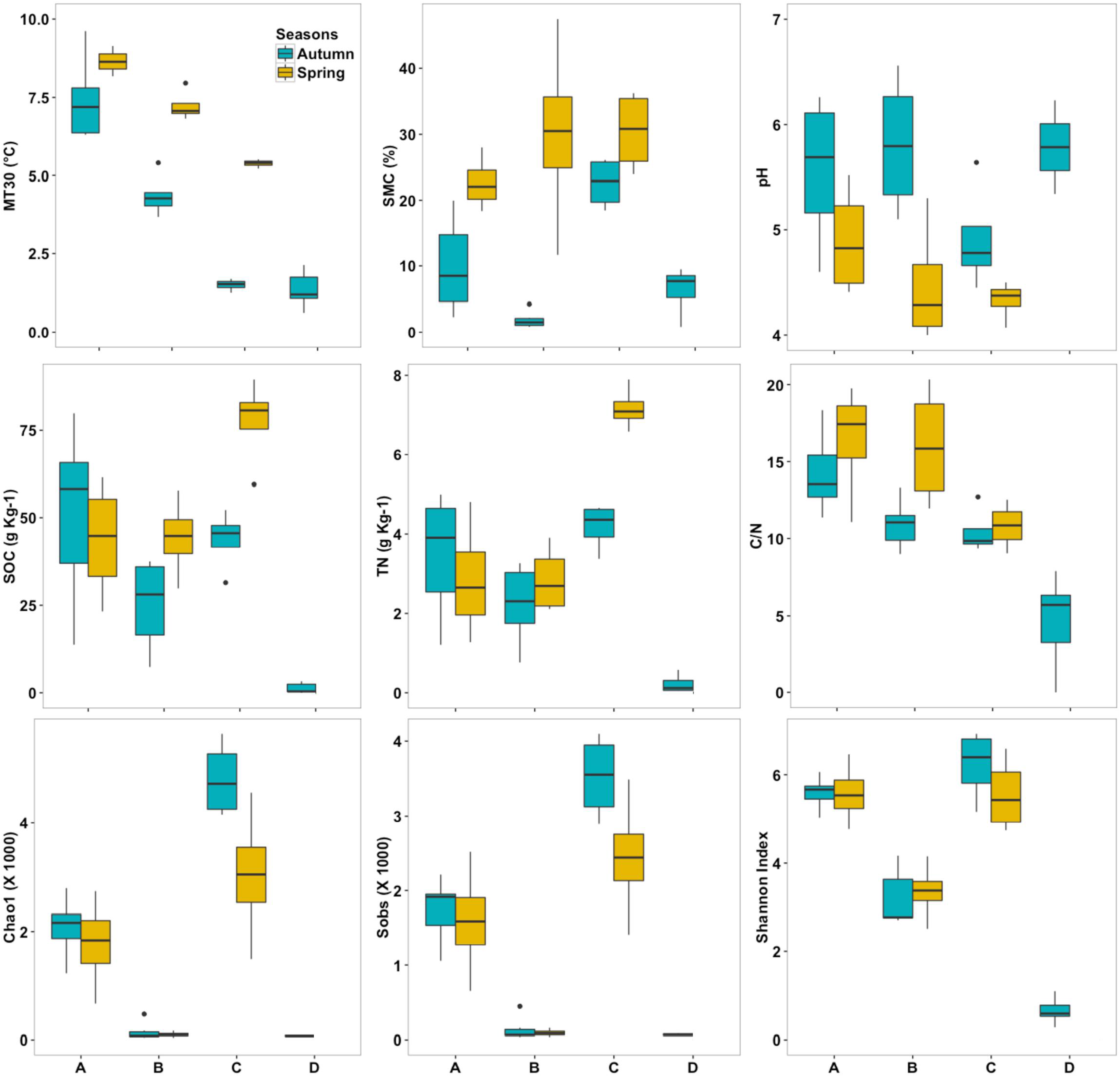
Environmental variables and alpha diversity indices for the four habitats along elevation gradient and seasons. Site abbreviations: A-Subalpine forest, B-Alpine scrub, CAlpine meadow and D-Morainic slope.

**Table 2:** Comparison of environmental variables and alpha diversity between habitats along elevation.

| Environmental variables |  |  |  |  |  |  |
| --- | --- | --- | --- | --- | --- | --- |
| Habitats | MT<sub>30</sub> (°C) |  | SMC (%) |  | pH |  |
| SF (n=11) | 8.01 ± 1.46 |  | 14.5 ± 8.62 |  | 5.33 ± 0.67 |  |
| AS (n=10) | 5.53 ± 1.57 |  | 13.1 ± 16.8 |  | 5.27 ± 0.89 |  |
| AM (n=8) | 3.43 ± 2.07 |  | 26.5 ± 6.34 |  | 4.62 ± 0.47 |  |
| MS (n=5) | 1.35 ± 0.59 |  | 6.34 ± 3.47 |  | 6.65 ± 0.85 |  |
| Comparison pairs | *p*-value | Cohen's *d* | *p*-value | Cohen's *d* | *p*-value | Cohen's *d* |
| SF vs. AS | **0.007** | 1.65 | 0.995 | 0.107 | 0.998 | 0.078 |
| SF vs. AM | **0.001** | 2.62 | **0.014** | 1.53 | 0.066 | 0.88 |
| SF vs. MS | **<0.001** | 5.21 | 0.072 | 1.09 | 0.076 | 1.18 |
| AS vs. AM | 0.143 | 1.13 | 0.149 | 1.73 | 0.239 | 0.88 |
| AS vs. MS | **<0.001** | 3.06 | 0.626 | 0.91 | 0.072 | 1.57 |
| AM vs. MS | 0.101 | 1.22 | **<0.001** | 3.67 | **0.013** | 3.18 |

| Habitats | SOC (g Kg<sup>-1</sup>) |  | TN (g Kg<sup>-1</sup>) |  | C/N ratio |  |
| --- | --- | --- | --- | --- | --- | --- |
| SF (n=11) | 48.2 ± 22.1 |  | 3.26 ± 1.51 |  | 15 ± 3 |  |
| AS (n=10) | 32.9 ± 15.1 |  | 2.48 ± 0.92 |  | 12.9 ± 3.65 |  |
| AM (n=8) | 60.6 ± 20.7 |  | 5.67 ± 1.67 |  | 10.6 ± 1.42 |  |
| MS (n=5) | 1.32 ± 1.47 |  | 0.22 ± 0.22 |  | 4.63 ± 3.08 |  |
| Comparison pairs | *p*-value | Cohen's *d* | *p*-value | Cohen's *d* | *p*-value | Cohen's *d* |
| SF vs. AS | 0.277 | 0.88 | 0.5 | 0.61 | 0.51 | 0.62 |
| SF vs. AM | 0.609 | 0.57 | **0.027** | 1.5 | **0.003** | 1.77 |
| SF vs. MS | **<0.001** | 2.5 | **<0.001** | 2.3 | **0.001** | 3.4 |
| AS vs. AM | **0.034** | 1.6 | **0.003** | 2.3 | 0.3 | 0.8 |
| AS vs. MS | **<0.001** | 2.5 | **<0.001** | 2.9 | **0.005** | 2.3 |
| AM vs. MS | **<0.001** | 3.5 | **<0.001** | 4.1 | **0.033** | 2.7 |

| Alpha diversity |  |  |  |  |  |  |
| --- | --- | --- | --- | --- | --- | --- |
| Habitats | Chao1 |  | Sobs |  | Shannon index (H') |  |
| SF (n=11) | 1967 ± 630 |  | 1682 ± 521 |  | 5.58 ± 0.45 |  |
| AS (n=10) | 128 ± 129 |  | 121 ± 122 |  | 3.25 ± 0.63 |  |
| AM (n=8) | 3921 ± 1340 |  | 2983 ± 883 |  | 5.88 ± 0.84 |  |
| MS (n=5) | 73 ± 11 |  | 69 ± 11 |  | 0.65 ± 0.35 |  |
| Comparison pairs | *p*-value | Cohen's *d* | *p*-value | Cohen's *d* | *p*-value | Cohen's *d* |
| SF vs. AS | **<0.001** | 3.94 | **<0.001** | 4.02 | **<0.001** | 4.25 |
| SF vs. AM | **0.005** | 1.98 | **0.012** | 1.8 | 0.525 | 0.46 |
| SF vs. MS | **<0.001** | 3.55 | **<0.001** | 3.66 | **<0.001** | 11.7 |
| AS vs. AM | **<0.001** | 4.25 | **<0.001** | 4.84 | **0.004** | 3.5 |
| AS vs. MS | 0.573 | 0.51 | 0.57 | 0.51 | **<0.001** | 4.7 |
| AM vs. MS | **<0.001** | 3.59 | **<0.001** | 4.1 | **0.001** | 7.5 |

Bold *p*-values indicate significant differences ( $p < 0.05$ ).
MT<sub>30</sub>- mean temperature of 30 days period; SMC- soil moisture content; SOC- soil organic carbon, TN- total nitrogen; C/N- carbon nitrogen ratio; Sobs- observed bacterial richness; Chao1- expected bacterial richness; Shannon index (H')- bacterial diversity.
Habitat abbreviations: SF- Subalpine forest; AS- Alpine scrub; AM- Alpine meadow; MS- Morainic slope

MT_30_ ranged from 1.3±0.59-8.01±1.46°C and was significantly different among all habitats, except AS vs AM and AM vs MS (Table 2). SMC was highest in AM (Mean=26.5%, SD=6.34, n=8), followed by SF (Mean=14.5%, SD=8.62, n=11), AS (Mean=13.1%, SD=16.8, n=10) and MS (Mean=6.34%, SD=3.47, n=5) (Figure 2, Table 2). AM had significantly higher moisture than SF and MS. We found no significant differences in moisture among SF vs AS, SF vs MS, AS vs AM and AS vs MS (Table 2).

Three of the habitats (AM, AS and SF) showed acidic pH, whereas MS showed near-neutral pH (Mean=6.65, SD=0.85, n=5) (Figure 2, Table 2). Overall, AM (Mean=4.62, SD=0.47, n=8) had lowest pH followed by SF (Mean=5.33, SD=0.67, n=11) and AS (Mean=5.27, SD=0.89, n=10). The soil pH of AM was significantly lower than MS. We found no significant difference in pH among all other habitats.

SOC and TN in MS was significantly lower than others and in AS was lower than AM, whereas all other habitats showed no significant difference except TN in SF was significantly lower than AM (Figure 2, Table 2). C:N ratio in all habitats was found to be below 20:1 (Figure 2, Table 2). MS showed significantly lower C:N value compared to other habitats. Further, AM showed significantly low values than SF.

### Seasonal differences in environmental variables

Between autumn and spring, AM and AS differed in both non-resource and resource variables, whereas SF varied in non-resource variables (Table 3). MT_30_ was significantly higher during spring in AM and AS (Figure 2, Table 3). Soil pH was significantly low in spring for AM and AS. SMC value showed significant increase in all habitats in spring. AM showed significantly high SOC and TN values in spring (Table 3). C:N ratio in AS was significantly higher during spring, but no differences were found in AM and SF (Table 3).

**Table 3:** Seasonal comparison of environmental variables and alpha diversity within habitats during autumn (October 2016) and spring (May 2017).

| Habitats |  | Autumn<br>(October 2016) | Spring<br>(May 2017) | *U* value | *p*-value | Cohen's *d* |
| --- | --- | --- | --- | --- | --- | --- |
| SF | **Environmental variables** |  |  |  |  |  |
|  | MT<sub>30</sub> (°C) | 7.35 ± 1.2 | 6.59 ± 1.17 | 5 | 0.315 | 0.64 |
|  | SMC (%) | 9.93 ± 6.8 | 22.6 ± 4.13 | 1 | **0.012** | 2.23 |
|  | pH | 5.58 ± 0.7 | 4.89 ± 0.52 | 5 | 0.109 | 1.17 |
|  | SOC (g Kg<sup>-1</sup>) | 50.9 ± 25 | 43.5 ± 17.1 | 11 | 0.648 | 0.34 |
|  | TN (g Kg<sup>-1</sup>) | 3.41 ± 1.5 | 2.8 ± 1.51 | 9 | 0.648 | 0.39 |
|  | C/N | 14.2 ± 2.4 | 16.42 ± 3.8 | 9 | 0.412 | 0.7 |
|  | **Alpha Diversity** |  |  |  |  |  |
|  | Sobs | 1736 ± 387 | 1587 ± 766 | 11.5 | 0.788 | 0.24 |
|  | Chao1 | 2079 ± 498 | 1769 ± 863 | 21 | 0.527 | 0.43 |
|  | Shannon index (H') | 5.59 ± 0.31 | 5.58 ± 0.7 | 12 | 0.788 | 0.01 |
| AS | **Environmental variables** |  |  |  |  |  |
|  | MT<sub>30</sub> (°C) | 4.3 ± 0.59 | 7.2 ± 0.49 | 0 | **0.01** | 5.2 |
|  | SMC (%) | 1.87 ± 1.27 | 30.0 ± 14.6 | 0 | **0.01** | 3.12 |
|  | pH | 5.81 ± 0.59 | 4.5 ± 0.58 | 2 | **0.038** | 2.3 |
|  | SOC (g Kg<sup>-1</sup>) | 25.3 ± 12.7 | 44.2 ± 11.5 | 3 | 0.067 | 1.56 |
|  | TN (g Kg<sup>-1</sup>) | 2.2 ± 0.9 | 2.8 ± 0.8 | 7 | 0.352 | 0.7 |
|  | C/N | 10.9 ± 1.53 | 15.9 ± 3.93 | 1 | **0.019** | 1.67 |
|  | **Alpha Diversity** |  |  |  |  |  |
|  | Sobs | 139 ± 156 | 93 ± 51 | 18 | 1 | 0.39 |
|  | Chao1 | 147 ± 165 | 99 ± 52 | 12 | 1 | 0.39 |
|  | Shannon index (H') | 3.1 ± 0.6 | 3.35 ± 0.66 | 17 | 1 | 0.39 |
| AM | **Environmental variables** |  |  |  |  |  |
|  | MT<sub>30</sub> (°C) | 1.49 ± 0.18 | 5.37 ± 0.13 | 0 | **0.029** | 24.7 |
|  | SMC (%) | 22.56 ± 8.59 | 30.4 ± 6.15 | 2 | **0.05** | 1.04 |
|  | pH | 4.91 ± 0.51 | 4.33 ± 0.18 | 1 | **0.05** | 1.52 |
|  | SOC (g Kg<sup>-1</sup>) | 43.7 ± 8.7 | 77.5 ± 12.7 | 0 | **0.029** | 3.1 |
|  | TN (g Kg<sup>-1</sup>) | 4.2 ± 0.6 | 7.1 ± 0.5 | 0 | **0.029** | 5.4 |
|  | C/N | 10.4 ± 1.53 | 10.8 ± 1.51 | 7 | 1 | 0.26 |
|  | **Alpha Diversity** |  |  |  |  |  |
|  | Sobs | 3522 ± 568 | 2444 ± 849 | 2 | 0.114 | 1.49 |
|  | Chao1 | 4807 ± 706 | 3034 ± 1263 | 2 | 0.114 | 1.73 |
|  | Shannon index (H') | 6.22 ± 0.8 | 5.55 ± 0.84 | 3 | 0.2 | 0.877 |

Bold *p*-values indicate significant differences (*p* < 0.05).
MT<sub>30</sub>- mean temperature of 30 days period; SMC- soil moisture content; SOC- soil organic carbon, TN- total nitrogen; C/N- carbon nitrogen ratio; Sobs- observed bacterial richness; Chao1- expected bacterial richness; Shannon index (H')- bacterial diversity.
Habitat abbreviations: SF- Subalpine forest; AS- Alpine scrub; AM- Alpine meadow; MS- Morainic slope

### Variation in alpha diversity along elevation and seasons

We found no specific elevation trend in bacterial richness (Chao1 and Sobs) but alpha diversity (H’) showed overall decreasing elevation trend, except highest diversity at AM (Figure 2, Table 2). Results indicate that except AS and MS, other habitats had significantly different richness (Table 2). Similarly, except SF and AM, other habitats had significantly different bacterial diversity (Table 2). However, contrary to the patterns among the habitats along elevation, we found no significant seasonal variation in bacterial richness and diversity within AM, AS and SF (Figure 2, Table 3).

### Taxonomic overview of the sequence data

We generated total 68,50,909 paired-end raw sequences from 36 soil samples. After quality controls, a total of 16,56,540 high quality sequences were obtained averaging to 46015±13616 sequence reads/sample (Supplementary Table S2). The rarefaction curves gradually flattened with increasing number of sequences for all samples (Supplementary Figure S3), indicating adequate sequencing depth and representation of the bacterial communities. Rarefaction at 23800 reads provided >90% Good’s coverage (Supplementary Table S2) across 34 samples. Remaining two samples were eliminated from downstream analyses. Overall, we identified 20943 OTUs from these samples (n=34), classified into 33 phyla (97.1% of the OTUs), 99 classes (87.1%), 252 orders (84%), 423 families (80%), 956 genera (75.4%) and 20495 species (1.2%). Due to low classification percentage at species level, we decided to conduct all further diversity analyses at phyla, class and genera level.

Of all the identified phyla, 12 were abundant in these habitats and accounted for 98% of the total bacterial sequences (Figure 3a). *Proteobacteria* (36.8%), *Actinobacteria* (28.38%) and *Firmicutes* (10.7%) were present across all habitats, whereas the remaining nine phyla (Supplementary Table S3) were found in at least one of the habitats (Figure 3a). The details of other abundant and rare phyla are provided in Figure 3a and 3b and Supplementary Table S3 and S4. Within *Proteobacteria* phyla, the abundant classes were *Gammaproteobacteria* (20.6%) and *Alphaproteobacteria* (15.32%), whereas *Actinobacteria* phyla had *Actinobacteria* (22.9%) and *Thermoleophilia* (4.25%) as abundant classes. Phyla *Firmicutes* had *Bacilli* (11.2%) as abundant class (Supplementary Table S5). Other abundant and rare bacterial classes are listed in supplementary Table S5 and S6. The most abundant bacterial genus was *Mycobacterium* (14.4%) (Supplementary Table S7). Other abundant and rare bacterial genera are listed in supplementary Table S7 and S8.

**Figure 3:**
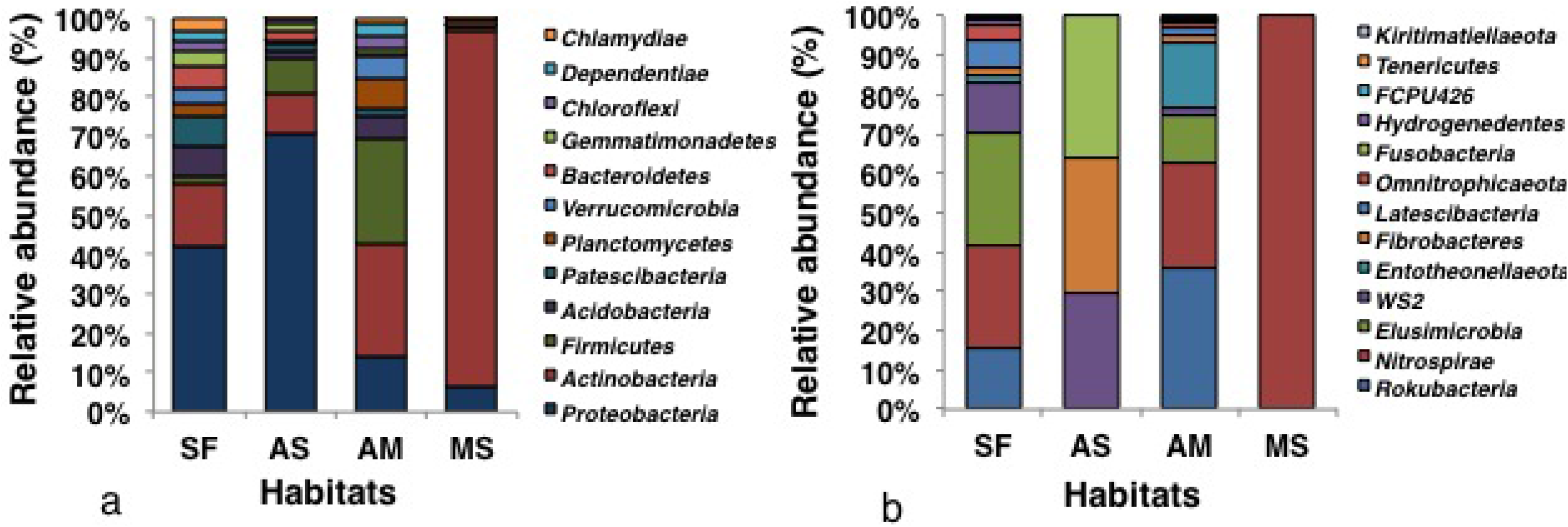
Relative abundance of (a) abundant and (b) rare bacterial phyla present in four habitats. Site abbreviations: MS-Morainic soils, AM-Alpine meadow, AS-Alpine scrub, SF-Subalpine forest.

### Habitat-wise comparisons in beta diversity

Three independent MRPP analysis using all, abundant and rare genera indicate significantly different community composition in the sampled habitats (A value ranged from 0.1-0.46, *p* < 0.01, Supplementary Table S9). Except no difference in rare genera of AS and MS all other pairwise MRPP analyses showed different community composition (A value ranged from 0.12-0.6, *p* < 0.05, Supplementary Table 9). The PCoA analyses showed a distinct MS cluster for all and abundant genera from other three habitats (Figure 4a-b), whereas MS and AS showed overlapping clusters for rare genera (similar to MRPP analyses) (Figure 4c). AM, AS and SF showed three distinct clusters indicating different community compositions for all three genera groups (Figure 4d-f). Combining both we interpret that community composition of all four habitats are different from each other.

**Figure 4:**
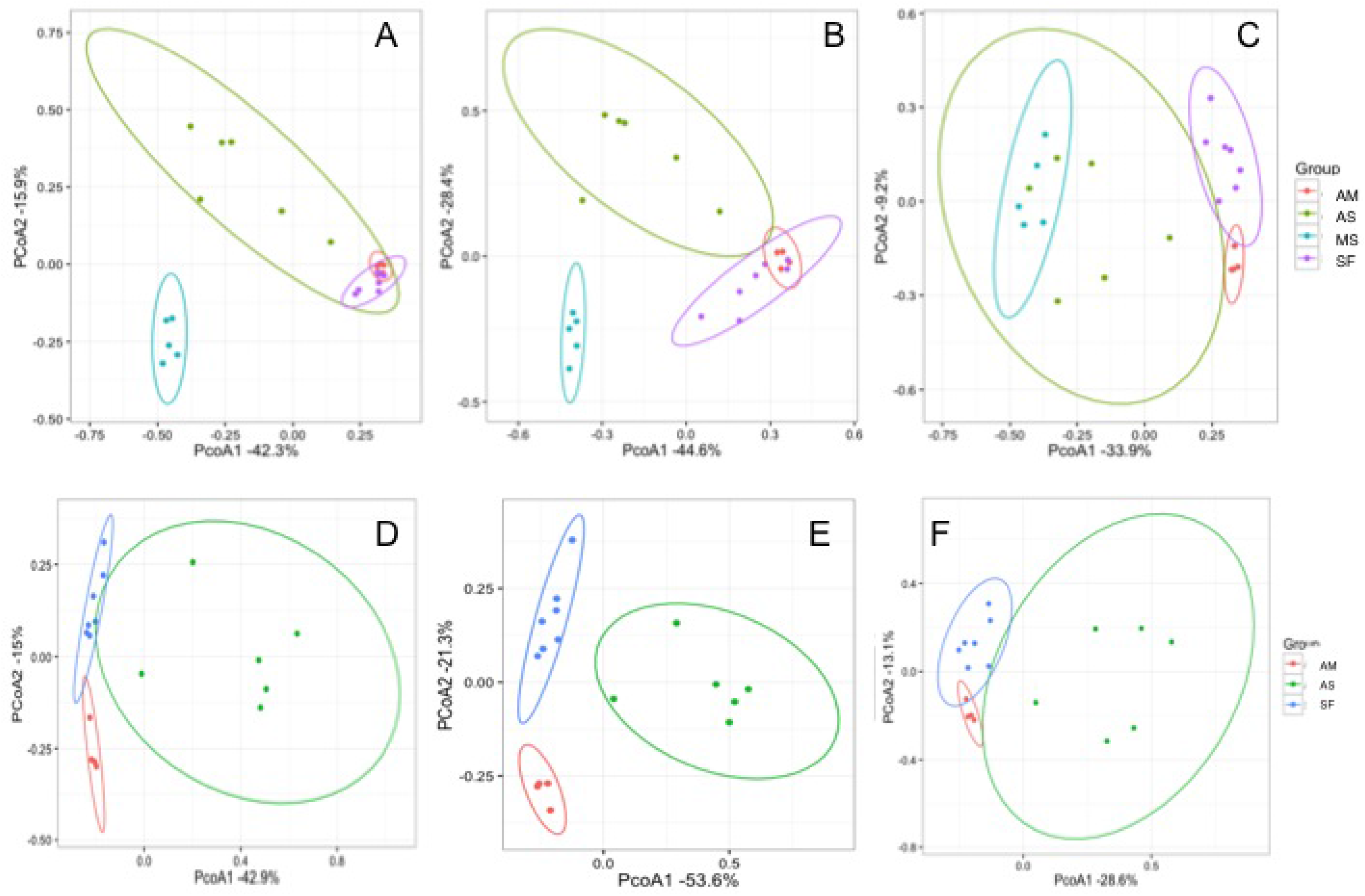
Principal co-ordinates analysis (PCoA) based on Bray-Curtis dissimilarity of bacterial communities for (a) all (b) abundant and (c) rare genera for four habitats and (d) all (e) abundant and (f) rare genera for three habitats. Site abbreviations: MS-Morainic soils, AM-Alpine meadow, AS-Alpine scrub, SF-Ssubalpine forest.

Our results indicate that variation in bacterial taxa composition and their relative abundances led to the observed differences in habitat wise community compositions along elevation. The average dissimilarities among the habitats ranged between 30-44% at phyla level (11 abundant and 8 rare phyla), 34-48% at class level (15 abundant and 64 rare class), and 49-69% at genera level (19 abundant and 318 rare genera) (Supplementary Table S10). In SF relative abundance of phyla *Bacteroidetes* (5.84%) and *Patescibacteria* (7.94%) were highest, whereas in AS *Proteobacteria* (69.73%) was highly abundant. In AM the phyla *Firmicutes* (26.6%), *Planctomycetes* (7.29%) and *Chloroflexi* (3.26%) and in MS the phylum *Actinobacteria* (90.1%) were highest compared to other habitats. Within rare phyla *Omnitrophicaeota* (0.01%) in SF and *Nitrospirae* (0.08%) and *Elusimicrobia* (0.037%) were highest in AM (Supplementary Table S3 and S4).

At class level, both SF and AS had the highest relative abundance of *Alphaproteobacteria* (20.4% and 22.6%, respectively), while AS was dominated by *Gammaproteobacteria* (46.4%). In SF *Bacteroidia* (5.79%) and *Saccharimonadia* (6.93%) were highly abundant. In AM *Bacilli* (28.6%), *Thermoleophilia* (8.85%)*, Planctomycetacia* (4.9%) and KD4-96 (1%) were highest, while MS was dominated by *Actinobacteria* (90%) (Supplementary Table S5).

At genus level, SF had the highest relative abundance of *Aquicella* (3.47%) and *Luteibacter* (1%), Ellin6055 (1.38%), *Flavobacterium* (0.94%) and *Mucilaginibacter* (0.83%). In AS, *Burkholderia-Caballeronia-Paraburkholderia* (16.7%)*, Acinetobacter* (12.5%)*, Pseudomonas* (3.8%), *Brucella* (8%) and *Sphingomonas* (4.6%) were highly abundant. *Nitrobacter* (0.68%), *Pseudonocardia* (2.96%)*, Nocardioides* (1.75%)*, Acidothermus* (1.57%) and *Solirubrobacter* (1.42%), *Bacillus* (4.7%) and *Sporosarcina* (3.67%) were highest in AM. The MS was dominated by *Mycobacterium* (89.3%) (Supplementary Table S6).

### Seasonal variation in beta diversity

The community composition in spring and autumn showed no significant variation within SF, AS and AM for all three genera groups (A value range −0.009-0.03, *p* > 0.05, Supplementary Table S11). Similarly, the PCoA analyses showed overlapping clusters for both seasons within the habitats (Supplementary Figure S4). Overall, no significant seasonal variation in the relative abundances of the abundant and rare taxonomic groups was found, with some exceptions in SF and AM. The relative abundance of abundant genera *Aquicella* and *Mucilaginibacter* decreased during spring in SF. Similarly, abundant genera *Allorhizobium* and *Nitrobacter* also decreased significantly in AM during spring (Supplementary Table S7).

### Relationship between environmental variables and bacterial richness, diversity and community composition along elevation

The results of Pearson correlation showed that bacterial richness (Sobs) was significantly correlated with SMC, SOC and TN, whereas bacterial diversity (Shannon index, H’) was significantly correlated to elevation and all other environmental variables (Supplementary Figure S2). Multiple regression analysis showed that bacterial richness was positively related with TN and SOC whereas diversity was related with MT_30_, TN and SOC (Table 4).

**Table 4:** Results of Multiple ordinary least square regression representing relationship of bacterial richness and diversity with environmental variables. The variation in richness and diversity is explained by only significant variables.

| Response variable | Environmental variables | R-squared | Estimate<sup>a</sup> | p-value<sup>b</sup> |
| --- | --- | --- | --- | --- |
| Bacterial richness (Sobs) | Intercept | 0.45 | 5.316 | 0.986794 |
|  | SMC |  | 1.912 | 0.90684 |
|  | TN |  | 398.428 | 0.000204 *** |
| Bacterial richness (Sobs) | Intercept | 0.32 | 119.401 | 0.74298 |
|  | SMC |  | 11.797 | 0.50975 |
|  | SOC |  | 24.82 | 0.00629 ** |
| Bacterial diversity (Shannon index) | Intercept | 0.55 | 1.472804 | 0.01349* |
|  | MT<sub>30</sub> |  | 0.193437 | 0.037753* |
|  | SMC |  | -0.004639 | 0.840405 |
|  | TN |  | 0.582613 | 0.000161*** |
| Bacterial diversity (Shannon index) | Intercept | 0.5218 | 1.779587 | 0.003779** |
|  | MT<sub>30</sub> |  | 0.099953 | 0.327195 |
|  | SMC |  | 0.002851 | 0.902853 |
|  | SOC |  | 0.002851 | 0.000514*** |

SMC- soil moisture content; SOC- soil organic carbon, TN- total nitrogen; MT<sub>30</sub>- mean temperature of 30 days period
<sup>a</sup>Estimate: $\beta$ coefficient
<sup>b</sup>p-value (significance value): \*\*\*, $P < 0.001$ ; \*\*, $P < 0.01$ ; \*, $P < 0.05$

Mantel test showed that the community composition of all and abundant phyla (n=12), class (n=19) and genera (n=24) were significantly correlated to elevation and all other environmental variables, except soil SMC (Supplementary Table S12). However, rare taxa community composition showed no correlation with elevation. For rare phyla (n=14 most abundant phyla) the community composition was significantly correlated to pH, where as at class and genus level the rare taxa (n=20) community composition was significantly correlated to pH, SOC, TN and C:N ratio.

Redundancy analysis with 12 abundant phyla showed that MT_30_, SOC and TN could significantly explain 34% of the variation in their relative abundance (F_(3,30)_ = 5.17, *p* = 0.001; Figure 5). Univariate regression analyses showed that relative abundances of *Proteobacteria, Bacteroidetes, Patescibacteria* and *Chlamydiae* were positively associated and *Actinobacteria* was negatively associated with MT_30_ (Supplementary Table S13). *Acidobacteria, Planctomycetes, Firmicutes, Verrucomicrobia* and *Chloroflexi* were associated positively with TN and SOC (Supplementary Table S13). None of the environmental variables explained the rare phyla distributions in these habitats.

**Figure 5:**
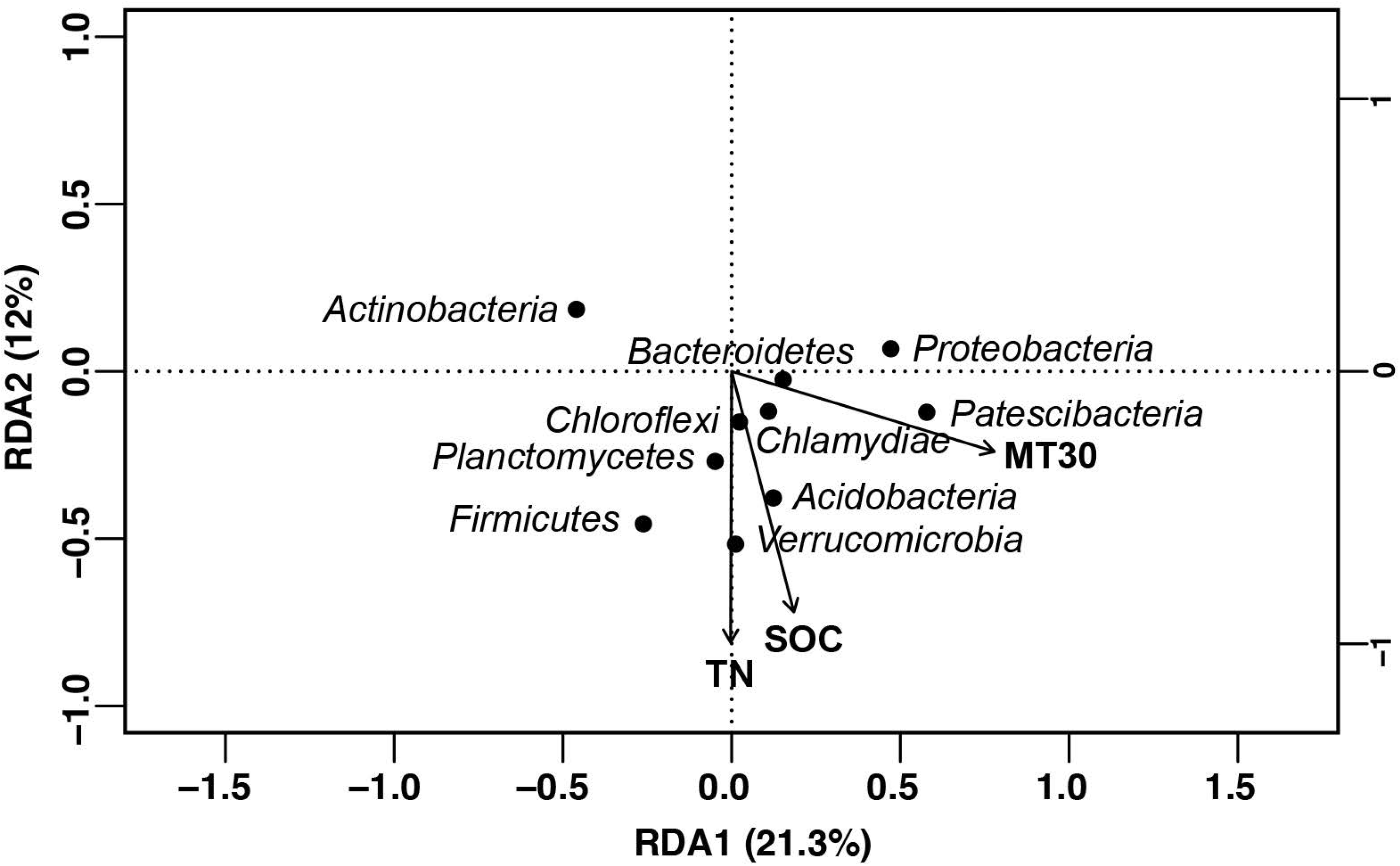
Redundancy analysis showing relation between bacterial phyla and environmental variables.

Similarly, at class level relative abundances of abundant (n=19) taxa was significantly influenced by MT_30_, SOC and TN (F_(3,30)_ = 5.24, *p* = 0.001). Univariate regression analyses showed that relative abundance of majority of the abundant bacterial class were significantly associated with the same environmental variable as the phyla they belong (Supplementary Table S14). While *Gammaproteobacteria* and *Alphaproteobacteria* were positively associated with MT_30_, *Deltaproteobacteria* showed positive relation with TN and SOC. Similarly, *Acidimicrobiia* and *Thermoleophilia* showed positive relation with TN and SOC. None of the environmental variables explained the rare class distributions in these habitats.

## Discussion

To date research efforts on soil bacterial community in the Himalayas have mostly focused on recording local diversity and communities from various habitats (cold desserts, glaciers, moraine lakes, surface snow etc.) [58, 59]. However, our knowledge regarding environmental drivers behind spatial and temporal patterns in bacterial communities is lacking within the Indian Himalayan region. To the best of our knowledge, this is the first study to address this gap where we identified the environmental factors influencing soil bacterial community alpha diversity and composition along elevation gradient covering four different alpine habitats across two seasons in the western Himalaya. Our results showed no elevation trend in the bacterial richness and a non-monotonous decrease in their diversity. This is in agreement with other studies reporting no clear microbial elevation pattern [60–62]. Unlike plant [63–65], animal and birds [66, 67] bacterial communities are known to exhibit inconsistent relationship with elevation. However, bacterial community richness, diversity and composition varied significantly between habitats as described from other alpine habitats from Tibetan Plateau and mountains in China and European Alps [22, 68, 69]

The habitat-specific patterns of bacterial community suggest that other factors than elevation might play crucial role in shaping their community [60]. Elevation provides comprehensive changes in habitat types where various biotic and abiotic factors influence bacterial diversity and composition [69]. For example, we found an unexpected result of highest bacterial richness and diversity in AM (higher elevation) when compared to SF and AS (lower elevation habitats). This pattern is contrasting as higher elevations are expected to retain low bacterial richness and diversity due to environmental harshness [70]. However, our results indicate that high amounts of SOC and TN from limited decomposition and turnover rates (due to low temperature) may have supported such richness and diversity. On the other hand, low SOC and TN in sparsely vegetated MS at highest elevation might have resulted in lowest bacterial richness and diversity. Overall, our results suggest that SOC and TN strongly influence bacterial community richness and diversity. Such patterns have also been described earlier from elevation gradient studies from Mt. Changbai, China and Italian Alps [22, 62, 71]. Habitat-specific variations in plant composition and chemistry of plant residues (root exudates and litter) regulate belowground environmental conditions by influencing the soil abiotic factors (nutrient quality, quantity, pH and soil moisture content etc.), which in turn affect the diversity and composition of microbial communities [22]. SOC and TN influence soil bacterial community by regulating growth, metabolism and cell formation [72, 73] and thus are generally found to be positively correlated [71, 74]. In our study, five out of the 12 abundant bacterial phyla (*Acidobacteria*, *Planctomycetes, Firmicutes, Verrucomicrobia* and *Chloroflexi*) and corresponding class were positively correlated with SOC and TN, indicating that they are the major drivers of bacterial community in different habitats. Similar patterns have been reported from other areas also [23, 71]. Another important abiotic factor, soil pH showed a surprisingly contrasting pattern when compared with other studies [23, 62, 75, 76]. While soil pH is known as one of the strongest driver of bacterial communities we didn’t find any effect of it in our study, possibly due to narrow pH range and limited variation among sampling sites.

Despite no clear elevation trend in our study, MT_30_ showed surprisingly strong relationship with bacterial diversity and composition. Earlier studies have suggested that temperature causes variations in soil bacterial community in natural environments by regulating their growth and metabolism [77], and a range of climatic factors (mean annual temperature, mean annual daily max temperature, mean annual precipitation, monthly mean air temperature, monthly mean precipitation etc.) are good predictors of variations in soil bacterial taxonomic diversity and composition [61, 69, 78]. Such effects of temperature can also be explained by species/group specific life strategies. For example, our RDA and univariate regression analyses showed a direct relationship between MT_30_ and relative abundances of phyla *Proteobacteria, Bacteroidetes, Patescibacteria* and *Chlamydiae*. As these bacterial groups are known to express copiotrophic life strategies [79] their abundances decrease in cold and oligotrophic environments of high elevation habitats (AM and MS). Contrarily, relative abundance of oligotrophic *Actinobacteria* [80, 81] showed negative relationship with MT_30_, where they were abundant in cold and low-nutrient habitats of AM and MS. While such explanations using bacterial “copiotrophic-oligotrophic” life-strategy spectrum could be over-simplified, it is known that the bacterial groups of related life-strategies tend to respond similarly to their respective microenvironment conditions (temperature, nutrient types, nutrient availability etc.) [79]. Nevertheless, such patterns have important implications in bacterial community shifts and soil carbon cycles under global warming scenarios. For instance, increasing temperatures under the current global scenario could lead to a gradual shift in bacterial community from Oligotrophs to Copiotrophs, which in turn can affect soil carbon cycles by increasing SOC decompositions [82]. Finally, it is important to point out that the unexplained variations in diversity and composition of abundant and rare bacterial communities indicate influences from other factors that were not measured in this study.

Interestingly, we did not find any seasonal difference in bacterial richness and diversity within each habitat types (except some exceptions in SF and AM) despite significant differences in some of the environmental variables (see Table 3). Earlier study by Siles and Margesin (2017) has also reported similar pattern in alpine forest soils. Plausible explanations could be (a) relatively stable bacterial community over short-term changes in environmental variables and (b) highly heterogeneous bacterial community within a habitat that can survive seasonal environmental variations. However, reports from subnival alpine regions [21], natural grasslands [83], and temperate forest ecosystems [84] showed significant seasonal effects on microbial diversity and community composition.

In conclusion, we here demonstrated that the soil bacterial community diversity and composition varied significantly among habitats along elevation but were stable seasonally within each habitat in alpine region of the western Himalaya. This pattern is influenced by a range of abiotic (temperature) and biotic (habitat type, soil organic carbon, nitrogen) factors but not directly by elevation gradient. These results provide the first baseline information on bacterial communities of different habitats and their relationship with local environmental factors. In the context of the various climate change prediction situations from the Himalayan region these results can help in modeling accurate climate adaptation scenarios of bacterial niches and their resulting impacts on soil carbon cycles. We suggest future expansion of such studies in other parts of the Indian Himalayan region where extensive spatial and seasonal sampling will help in understanding local responses of soil bacterial communities to varying climatic conditions. Given the critical role played by the Himalaya in global carbon cycle such information will help in taking informed decisions to mitigate effects of climate warming in this sensitive region.

## Acknowledgements

We acknowledge the Forest Department of Uttarakhand for providing necessary permits to carry out the research. Our thanks to the Forest Department officials and frontline staffs for their support and assistance during field sampling. We acknowledge help from Dr. Devendra, Dr. Ishwari and Umed during field surveys. We appreciate technical help from Arun (laboratory work), Debanjan (GIS), Dr. Awadhesh and Tejali (NGS work), Sitendu, Nilanjan and Dr. Raman (Analysis). We thank the Director, Dean, Research Coordinator and Nodal Officer of Wildlife Forensics and Conservation Genetics Cell of Wildlife Institute of India and Dr. Sathyakumar, Nodal Scientist, NMSHE for their support in this work.

## Declarations

### Funding

This research is part of the project National Mission for Sustaining the Himalayan Ecosystem (NMSHE) Programme, funded by Department of Science and Technology, Government of India (Grant no. DST/SPLICE/CCP/NMSHE/TF-2/WII/2014[G]). Partial funding for the bacterial NGS work was supported by United Nations Development Programme and the Ministry of Environment, Forest and Climate Change Government of India through the Third National Communication project (7/2/2015-CC). Pamela Bhattacharya was supported by Council of Scientific and Industrial Research, Government of India (Award no. 09/668(0012)/2019-EMR-I) and Samrat Mondol was supported by the Department of Science and Technology INSPIRE Faculty Award (No.IFA12-LSBM-47).

### Competing Interests

The authors declare there are no competing interests.

### Ethics approval

Not applicable

### Consent to participate

Not applicable

### Consent for publication

Not applicable

### Availability of data and material

The data set has been deposited to National Centre for Biotechnology Information (NCBI) Short Read Archive (https://www.ncbi.nlm.nih.gov/sra) under BioProject accession number PRJNA658431.

### Code availability

Not applicable

### Authors’ contributions

GSR, GT and PB conceived the idea and design of the study. GSR and GT supervised fieldwork and data analysis. SM supervised laboratory work and data analysis. PB conducted field sampling, laboratory work and data analyses. PB and SM prepared the draft manuscript. All authors reviewed and approved the final version.

---

**Figure S1.**
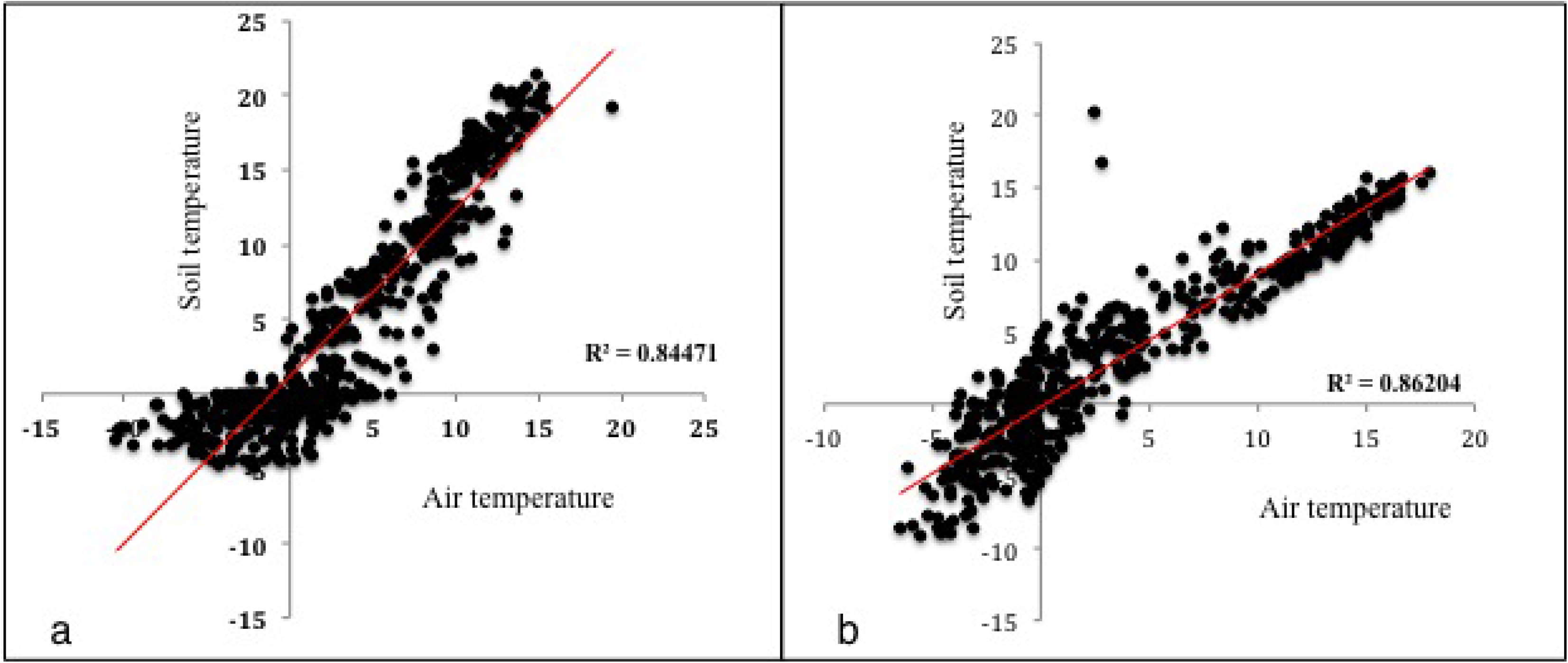
Scatter plot of air temperature (5 ft above ground) and soil temperature (5 cm below ground) at (a) 3564 m and (b) 4020 m elevation.

**Figure S2.**
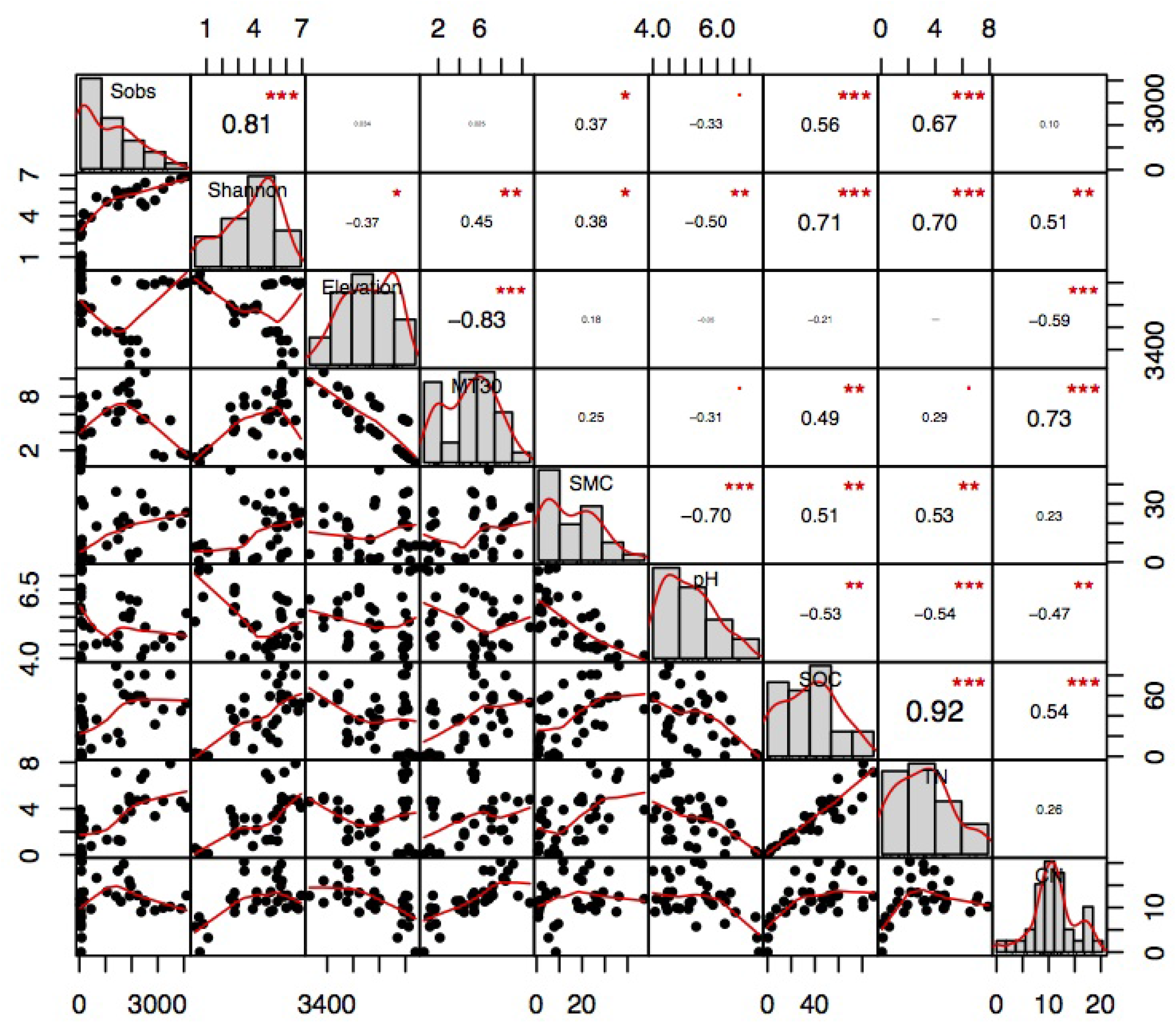
Pearson correlation analysis of bacterial richness, diversity and environmental variables.

**Figure S3:**
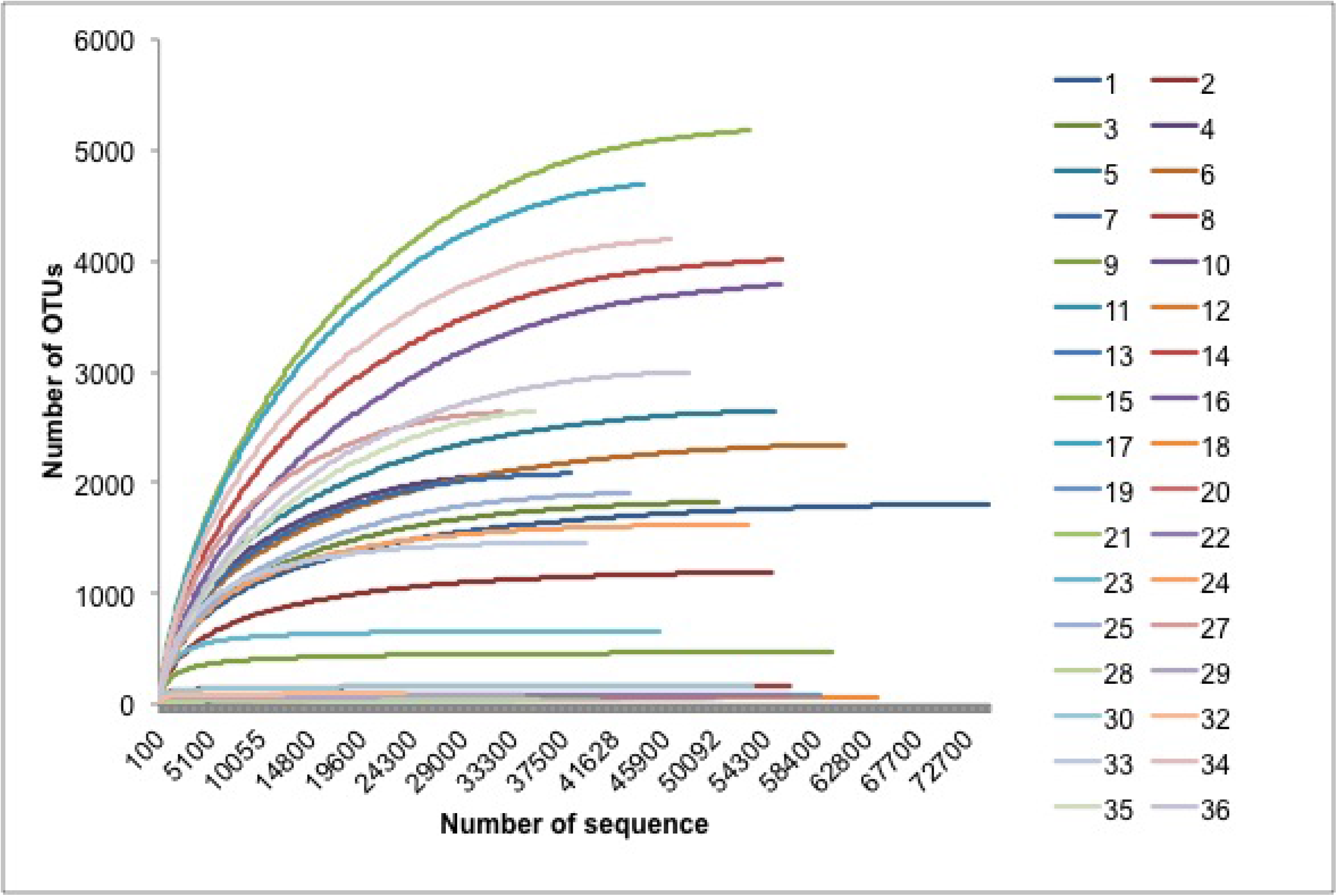
Rarefaction curve of samples (n=34) in this study representing expected number of operational taxonomic units (OTUs) for a given number of sequences per sample.

**Table S1:**
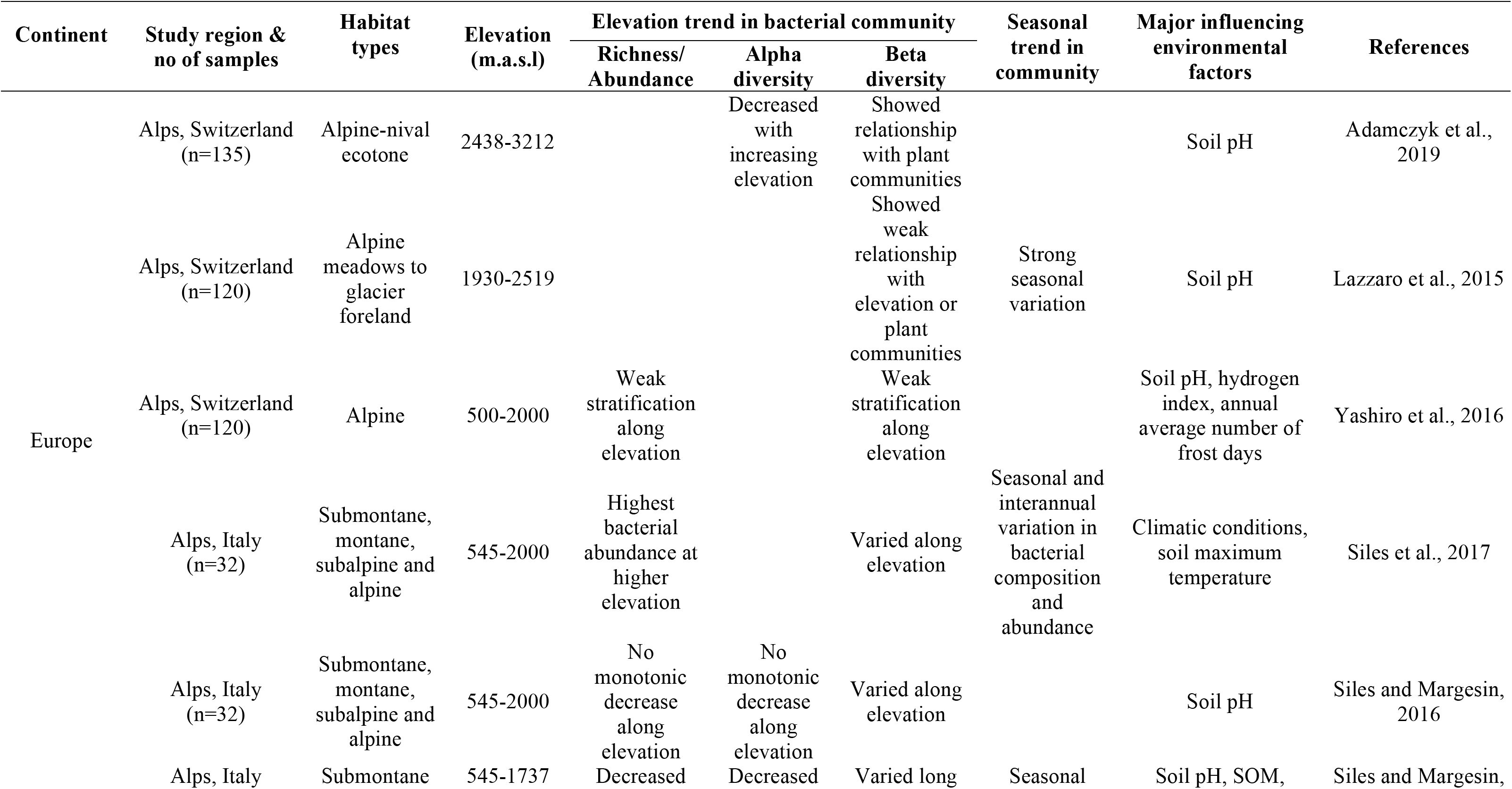
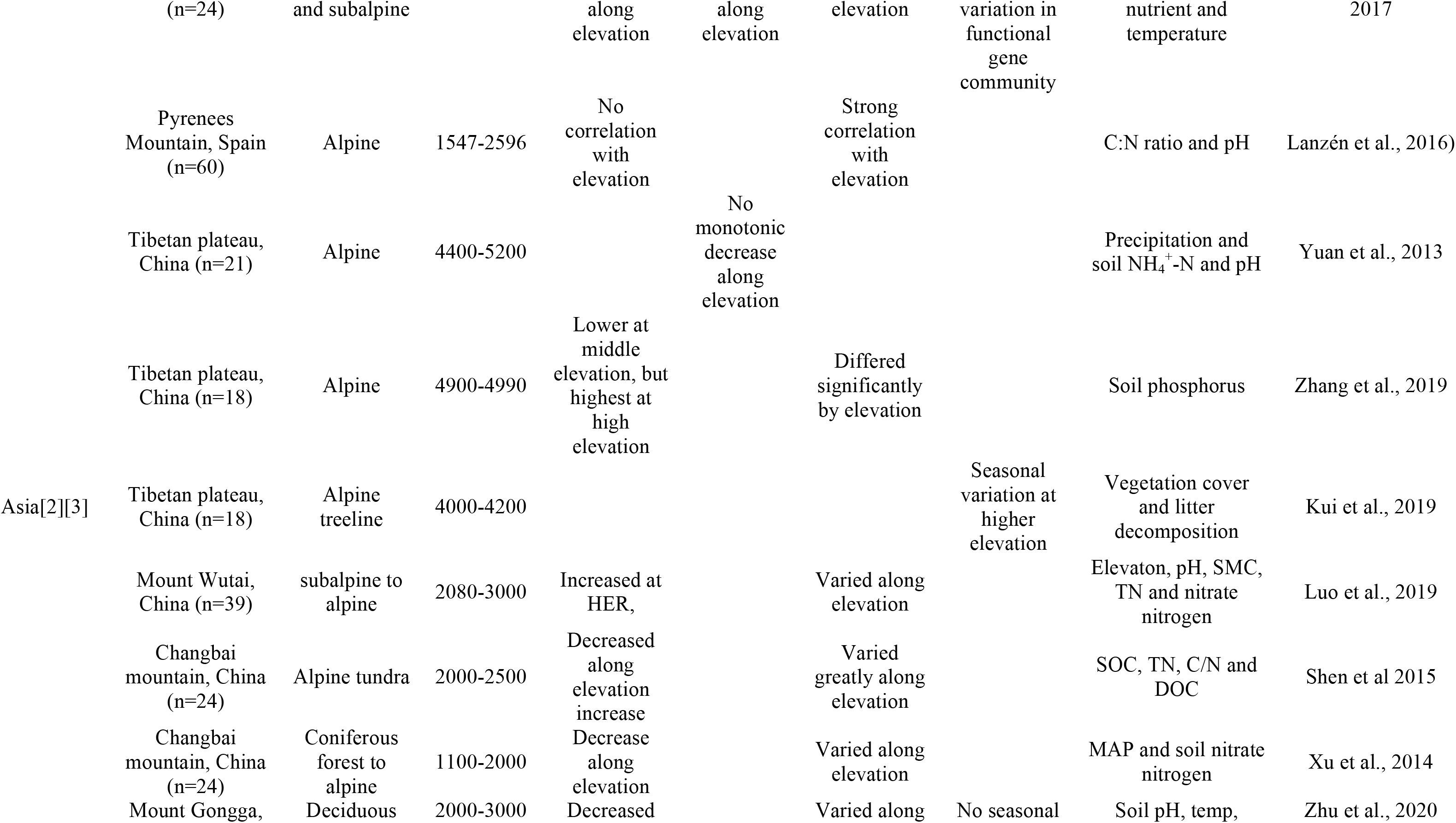
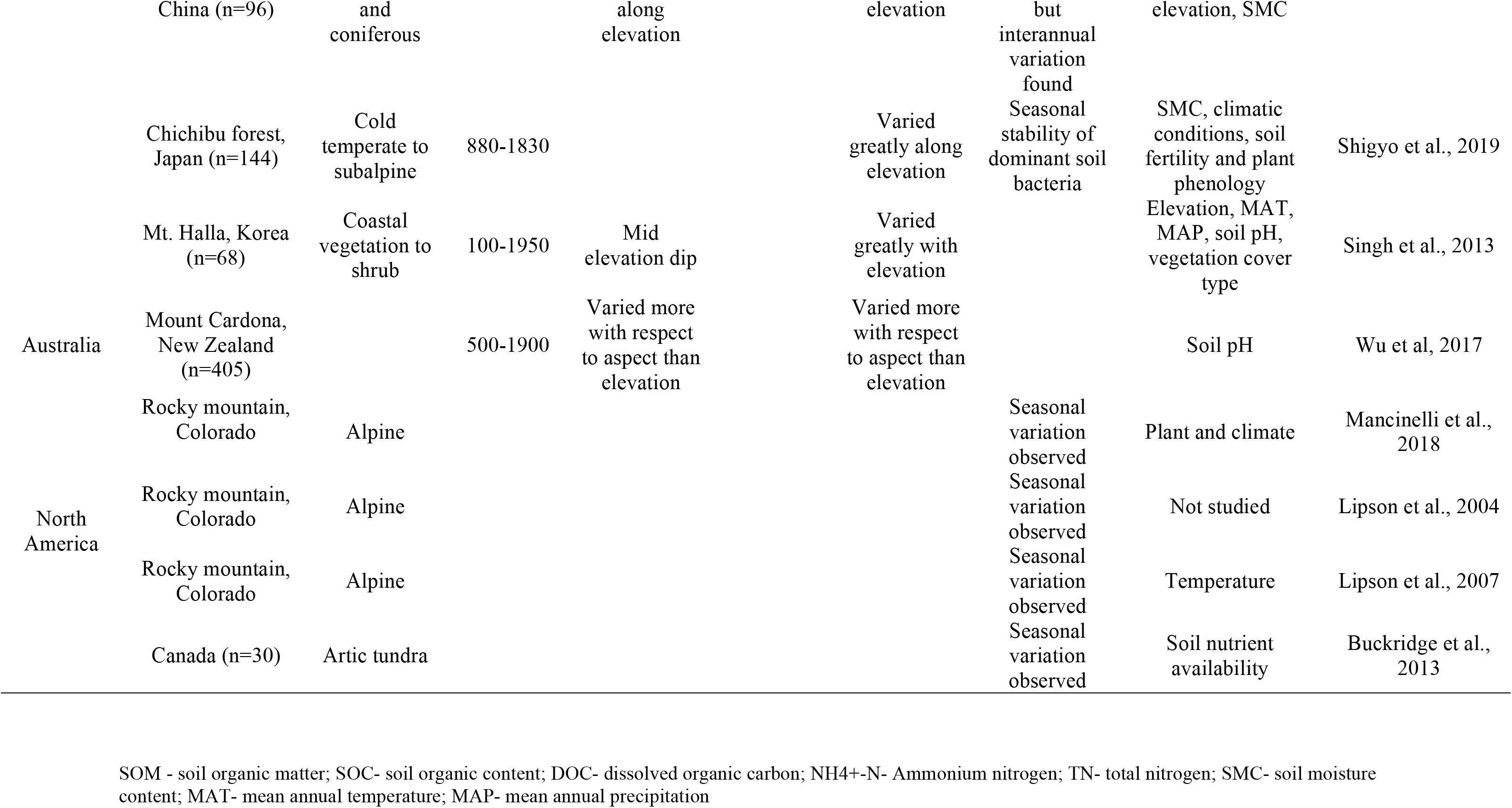
Continent-wise work done on soil bacterial community in alpine vegetation types along elevation and seasons.

**Table S2:** Number of sequences and Good’s coverage at all sampling sites.

| Sample No | Sample ID | Year of Sampling | Seasons | Total number of paired end reads | Reads after quality trimming and alignment | Good's coverage (%) |
| --- | --- | --- | --- | --- | --- | --- |
| 1 | SF1a | October-16 | Autumn | 292923.44 | 74824 | 98.2426 |
| 2 | SF1b | October-16 | Autumn | 186084.79 | 54884 | 98.9792 |
| 3 | SF2 | October-16 | Autumn | 207662.01 | 50310 | 98.1161 |
| 4 | SF3a | October-16 | Autumn | 181176.80 | 32326 | 98.0726 |
| 5 | SF3b | October-16 | Autumn | 262308.33 | 54936 | 96.9434 |
| 6 | SF4 | October-16 | Autumn | 244920.87 | 60880 | 97.3816 |
| 7 | SF5 | October-16 | Autumn | 164836.77 | 38126 | 97.9065 |
| 8 | AS1 | October-16 | Autumn | 177914.78 | 56168 | 99.9561 |
| 9 | AS2 | October-16 | Autumn | 185905.23 | 59984 | 99.8465 |
| 10 | AS3 | October-16 | Autumn | 146491.65 | 42202 | 99.982 |
| 11 | AS4 | October-16 | Autumn | 134401.22 | 33410 | 99.9825 |
| 12 | AS5a | October-16 | Autumn | 163819.26 | 49458 | 99.9801 |
| 13 | AS5b | October-16 | Autumn | 157684.27 | 50541 | 99.9865 |
| 14 | AM1 | October-16 | Autumn | 235374.74 | 56467 | 94.6964 |
| 15 | AM2 | October-16 | Autumn | 285827.61 | 53606 | 92.6876 |
| 16 | AM3 | October-16 | Autumn | 301118.98 | 56241 | 94.5346 |
| 17 | AM4 | October-16 | Autumn | 311183.64 | 46132 | 93.4249 |
| 18 | MS1 | October-16 | Autumn | 180338.85 | 64044 | 99.9692 |
| 19 | MS2 | October-16 | Autumn | 168068.86 | 58742 | 99.972 |
| 20 | MS3 | October-16 | Autumn | 144905.53 | 50092 | 99.9721 |
| 21 | MS4 | October-16 | Autumn | 164238.23 | 40601 | 99.9772 |
| 22 | MS5 | October-16 | Autumn | 163849.19 | 56245 | 99.9651 |
| 23 | SF1 | May-17 | Spring | 152866.05 | 45477 | 99.8848 |
| 24 | SF2 | May-17 | Spring | 197217.56 | 52823 | 98.7087 |
| 25 | SF3 | May-17 | Spring | 220650.24 | 42891 | 97.9068 |
| 26 | SF4 | May-17 | Spring | 187102.30 | 32600 | 97.6476 |
| 27 | SF5 | May-17 | Spring | 36450.83 | 10055 | Eliminated |
| 28 | AS1 | May-17 | Spring | 142750.80 | 35403 | 99.9904 |
| 29 | AS2 | May-17 | Spring | 146641.28 | 34210 | 99.9601 |
| 30 | AS3 | May-17 | Spring | 186443.91 | 53241 | 99.9546 |
| 31 | AS4 | May-17 | Spring | 127308.57 | 23804 | 99.9874 |
| 32 | AS5 | May-17 | Spring | 56771.12 | 12421 | Eliminated |
| 33 | AM1 | May-17 | Spring | 147000.40 | 39475 | 99.2639 |
| 34 | AM2 | May-17 | Spring | 294121.24 | 48029 | 94.3547 |
| 35 | AM3 | May-17 | Spring | 273905.53 | 36726 | 96.4695 |
| 36 | AM4 | May-17 | Spring | 220644.92 | 49166 | 96.0346 |

**Table S3:**
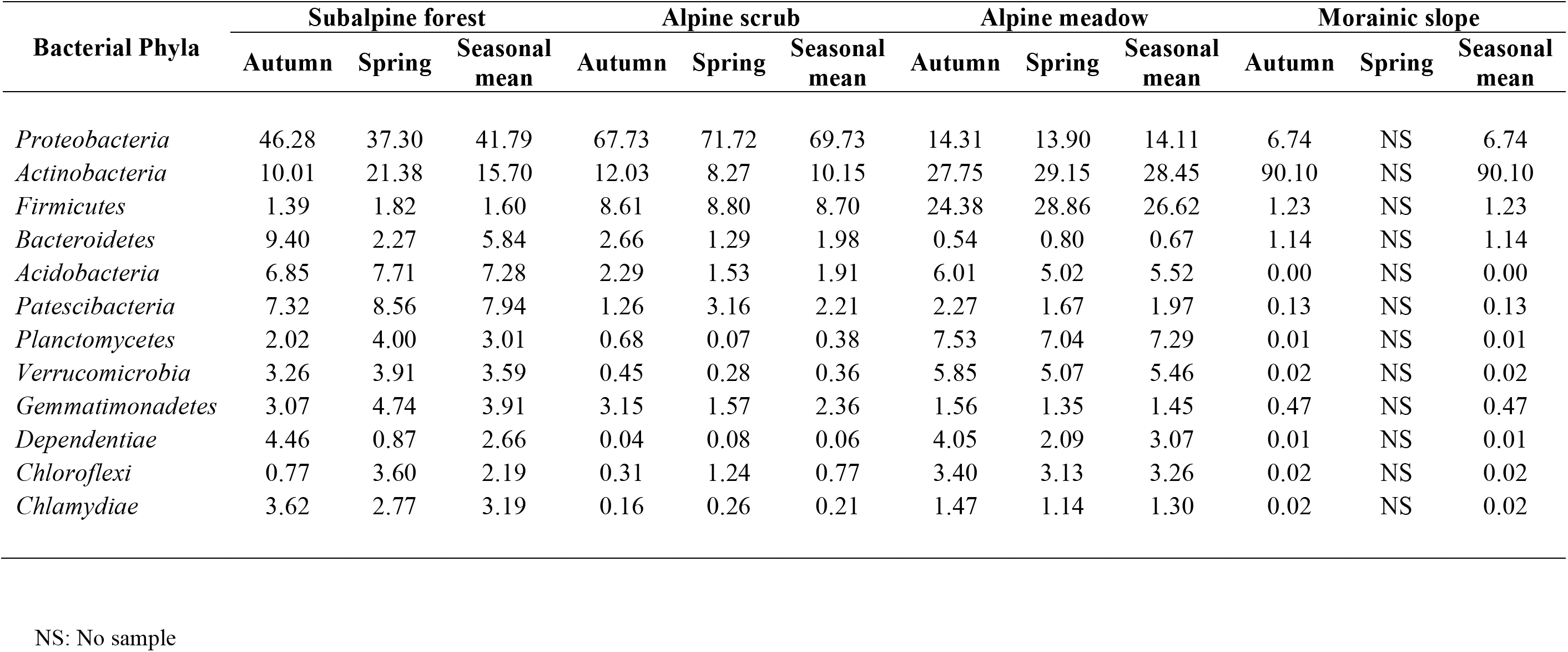
Relative abundance of abundant bacterial phyla (≥ 1% relative abundances) found in the four habitats in autumn and spring.

| Bacterial Phyla | Subalpine forest |  |  | Alpine scrub |  |  | Alpine meadow |  |  | Morainic slope |  |  |
| --- | --- | --- | --- | --- | --- | --- | --- | --- | --- | --- | --- | --- |
|  | Autumn | Spring | Seasonal mean | Autumn | Spring | Seasonal mean | Autumn | Spring | Seasonal mean | Autumn | Spring | Seasonal mean |
| *Proteobacteria* | 46.28 | 37.30 | 41.79 | 67.73 | 71.72 | 69.73 | 14.31 | 13.90 | 14.11 | 6.74 | NS | 6.74 |
| *Actinobacteria* | 10.01 | 21.38 | 15.70 | 12.03 | 8.27 | 10.15 | 27.75 | 29.15 | 28.45 | 90.10 | NS | 90.10 |
| *Firmicutes* | 1.39 | 1.82 | 1.60 | 8.61 | 8.80 | 8.70 | 24.38 | 28.86 | 26.62 | 1.23 | NS | 1.23 |
| *Bacteroidetes* | 9.40 | 2.27 | 5.84 | 2.66 | 1.29 | 1.98 | 0.54 | 0.80 | 0.67 | 1.14 | NS | 1.14 |
| *Acidobacteria* | 6.85 | 7.71 | 7.28 | 2.29 | 1.53 | 1.91 | 6.01 | 5.02 | 5.52 | 0.00 | NS | 0.00 |
| *Patescibacteria* | 7.32 | 8.56 | 7.94 | 1.26 | 3.16 | 2.21 | 2.27 | 1.67 | 1.97 | 0.13 | NS | 0.13 |
| *Planctomycetes* | 2.02 | 4.00 | 3.01 | 0.68 | 0.07 | 0.38 | 7.53 | 7.04 | 7.29 | 0.01 | NS | 0.01 |
| *Verrucomicrobia* | 3.26 | 3.91 | 3.59 | 0.45 | 0.28 | 0.36 | 5.85 | 5.07 | 5.46 | 0.02 | NS | 0.02 |
| *Gemmatimonadetes* | 3.07 | 4.74 | 3.91 | 3.15 | 1.57 | 2.36 | 1.56 | 1.35 | 1.45 | 0.47 | NS | 0.47 |
| *Dependentiae* | 4.46 | 0.87 | 2.66 | 0.04 | 0.08 | 0.06 | 4.05 | 2.09 | 3.07 | 0.01 | NS | 0.01 |
| *Chloroflexi* | 0.77 | 3.60 | 2.19 | 0.31 | 1.24 | 0.77 | 3.40 | 3.13 | 3.26 | 0.02 | NS | 0.02 |
| *Chlamydiae* | 3.62 | 2.77 | 3.19 | 0.16 | 0.26 | 0.21 | 1.47 | 1.14 | 1.30 | 0.02 | NS | 0.02 |

NS: No sample

**Table S4:** Relative abundance of rare bacterial phyla (≤ 0.1% relative abundances) found in the four habitats in autumn and spring.

| Bacterial Phyla | Subalpine forest |  |  | Alpine scrub |  |  | Alpine meadow |  |  | Morainic slope |  |  |
| --- | --- | --- | --- | --- | --- | --- | --- | --- | --- | --- | --- | --- |
|  | Autumn | Spring | Seasonal mean | Autumn | Spring | Seasonal mean | Autumn | Spring | Seasonal mean | Autumn | Spring | Seasonal mean |
| *Rokubacteria* | 0.024 | 0.050 | 0.037 | 0.000 | 0.000 | 0.000 | 0.141 | 0.075 | 0.108 | 0.000 | NS | 0.000 |
| *Nitrospirae* | 0.015 | 0.105 | 0.060 | 0.000 | 0.000 | 0.000 | 0.086 | 0.074 | 0.080 | 0.003 | NS | 0.003 |
| *Elusimicrobia* | 0.024 | 0.111 | 0.068 | 0.000 | 0.000 | 0.000 | 0.042 | 0.032 | 0.037 | 0.000 | NS | 0.000 |
| WS2 | 0.038 | 0.021 | 0.030 | 0.018 | 0.000 | 0.009 | 0.007 | 0.004 | 0.006 | 0.000 | NS | 0.000 |
| *Entotheonellaeota* | 0.002 | 0.005 | 0.004 | 0.000 | 0.000 | 0.000 | 0.066 | 0.033 | 0.050 | 0.000 | NS | 0.000 |
| *Fibrobacteres* | 0.006 | 0.005 | 0.006 | 0.021 | 0.000 | 0.011 | 0.009 | 0.004 | 0.007 | 0.000 | NS | 0.000 |
| *Latescibacteria* | 0.003 | 0.027 | 0.015 | 0.000 | 0.000 | 0.000 | 0.009 | 0.004 | 0.007 | 0.000 | NS | 0.000 |
| *Omnitrophicaeota* | 0.010 | 0.009 | 0.010 | 0.000 | 0.000 | 0.000 | 0.003 | 0.001 | 0.002 | 0.000 | NS | 0.000 |
| *Fusobacteria* | 0.000 | 0.000 | 0.000 | 0.000 | 0.022 | 0.011 | 0.001 | 0.000 | 0.001 | 0.000 | NS | 0.000 |
| *Hydrogenedentes* | 0.003 | 0.003 | 0.003 | 0.000 | 0.000 | 0.000 | 0.000 | 0.003 | 0.002 | 0.000 | NS | 0.000 |
| FCPU426 | 0.003 | 0.001 | 0.002 | 0.000 | 0.000 | 0.000 | 0.001 | 0.000 | 0.001 | 0.000 | NS | 0.000 |
| *Tenericutes* | 0.000 | 0.000 | 0.000 | 0.000 | 0.000 | 0.000 | 0.002 | 0.002 | 0.002 | 0.000 | NS | 0.000 |
| *Kiritimatiellaeota* | 0.000 | 0.001 | 0.001 | 0.000 | 0.000 | 0.000 | 0.002 | 0.001 | 0.002 | 0.000 | NS | 0.000 |

NS: No sample

**Table S5:** Relative abundance of abundant bacterial class (≥ 1% relative abundances) found in the four habitats in autumn and spring.

| Bacterial phyla | Bacterial class | Subalpine forest |  |  | Alpine scrub |  |  | Alpine meadow |  |  | Morainic slope |  |  |
| --- | --- | --- | --- | --- | --- | --- | --- | --- | --- | --- | --- | --- | --- |
|  |  | Autumn | Spring | Seasonal mean | Autumn | Spring | Seasonal mean | Autumn | Spring | Seasonal mean | Autumn | Spring | Seasonal mean |
| *Proteobacteria* | *Gammaproteobacteria* | 23.30 | 17.10 | 20.20 | 42.60 | 50.20 | 46.40 | 2.71 | 3.45 | 3.08 | 5.27 | NS | 5.27 |
| *Proteobacteria* | *Alphaproteobacteria* | 22.40 | 18.40 | 20.40 | 24.50 | 20.80 | 22.65 | 10.50 | 9.29 | 9.90 | 1.38 | NS | 1.38 |
| *Proteobacteria* | *Deltaproteobacteria* | 0.51 | 1.57 | 1.04 | 0.54 | 0.61 | 0.58 | 1.10 | 0.71 | 0.91 | 0.03 | NS | 0.03 |
| *Actinobacteria* | *Actinobacteria* | 7.89 | 11.40 | 9.65 | 8.67 | 7.14 | 7.91 | 15.10 | 20.20 | 17.65 | 90.00 | NS | 90.00 |
| *Actinobacteria* | *Thermoleophilia* | 1.25 | 7.36 | 4.31 | 2.84 | 0.57 | 1.71 | 9.46 | 8.24 | 8.85 | 0.06 | NS | 0.06 |
| *Actinobacteria* | *Acidimicrobiia* | 0.64 | 1.83 | 1.24 | 0.34 | 0.17 | 0.26 | 1.64 | 1.25 | 1.45 | 0.00 | NS | 0.00 |
| *Firmicutes* | *Bacilli* | 1.40 | 1.61 | 1.51 | 8.54 | 8.58 | 8.56 | 24.10 | 33.20 | 28.65 | 1.14 | NS | 1.14 |
| *Patescibacteria* | *Saccharimonadia* | 6.33 | 7.53 | 6.93 | 1.02 | 2.82 | 1.92 | 1.28 | 0.80 | 1.04 | 0.10 | NS | 0.10 |
| *Bacteroidetes* | *Bacteroidia* | 9.36 | 2.22 | 5.79 | 2.65 | 1.30 | 1.98 | 0.52 | 0.98 | 0.75 | 1.15 | NS | 1.15 |
| *Verrucomicrobia* | *Verrucomicrobiae* | 3.20 | 3.94 | 3.57 | 0.43 | 0.26 | 0.35 | 5.92 | 4.23 | 5.08 | 0.01 | NS | 0.01 |
| *Acidobacteria* | *Subgroup_6* | 2.93 | 4.88 | 3.91 | 0.80 | 1.48 | 1.14 | 2.75 | 1.64 | 2.20 | 0.00 | NS | 0.00 |
| *Acidobacteria* | *Acidobacteriia* | 2.62 | 0.66 | 1.64 | 0.22 | 0.00 | 0.11 | 1.97 | 0.94 | 1.46 | 0.00 | NS | 0.00 |
| *Acidobacteria* | *Blastocatellia_Subgroup_4* | 0.86 | 1.00 | 0.93 | 0.18 | 0.00 | 0.09 | 0.89 | 1.00 | 0.95 | 0.00 | NS | 0.00 |
| *Planctomycetes* | *Planctomycetacia* | 1.50 | 3.16 | 2.33 | 0.62 | 0.06 | 0.34 | 5.50 | 4.45 | 4.98 | 0.00 | NS | 0.00 |
| *Planctomycetes* | *Phycisphaerae* | 0.46 | 0.66 | 0.56 | 0.04 | 0.00 | 0.02 | 1.98 | 1.99 | 1.99 | 0.00 | NS | 0.00 |
| *Chlamydiae* | *Chlamydiae* | 3.65 | 2.77 | 3.21 | 0.16 | 0.26 | 0.21 | 1.43 | 0.77 | 1.10 | 0.01 | NS | 0.01 |
| *Gemmatimonadetes* | *Gemmatimonadetes* | 2.82 | 4.64 | 3.73 | 3.10 | 1.52 | 2.31 | 1.46 | 1.14 | 1.30 | 0.46 | NS | 0.46 |
| *Dependentiae* | *Babeliae* | 4.41 | 0.86 | 2.64 | 0.04 | 0.09 | 0.07 | 4.13 | 0.14 | 2.14 | 0.00 | NS | 0.00 |
| *Chloroflexi* | *KD4-96* | 0.31 | 2.03 | 1.17 | 0.12 | 0.34 | 0.23 | 1.04 | 0.91 | 0.98 | 0.00 | NS | 0.00 |

NS: No sample

**Table S6:** Relative abundance of rare bacterial class (≤ 0.1% relative abundances) found in the four habitats in autumn and spring.

| Bacterial phyla | Bacterial class | Subalpine forest |  |  | Alpine scrub |  |  | Alpine meadow |  |  | Morainic slope |  |  |
| --- | --- | --- | --- | --- | --- | --- | --- | --- | --- | --- | --- | --- | --- |
|  |  | Autumn | Spring | Seasonal mean | Autumn | Spring | Seasonal mean | Autumn | Spring | Seasonal mean | Autumn | Spring | Seasonal mean |
| *Nitrospirae* | *Nitrospira* | 0.017 | 0.108 | 0.062 | 0.000 | 0.000 | 0.000 | 0.092 | 0.077 | 0.085 | 0.006 | NS | 0.006 |
| *Patescibacteria* | WS6_Dojkabacteria | 0.105 | 0.112 | 0.109 | 0.000 | 0.000 | 0.000 | 0.001 | 0.002 | 0.002 | 0.000 | NS | 0.000 |
| *Armatimonadetes* | *Chthonomonadetes* | 0.014 | 0.002 | 0.008 | 0.000 | 0.000 | 0.000 | 0.113 | 0.068 | 0.091 | 0.000 | NS | 0.000 |
| *Acidobacteria* | Subgroup_5 | 0.023 | 0.048 | 0.036 | 0.000 | 0.000 | 0.000 | 0.065 | 0.039 | 0.052 | 0.000 | NS | 0.000 |
| *Bacteroidetes* | *Ignavibacteria* | 0.115 | 0.045 | 0.080 | 0.000 | 0.000 | 0.000 | 0.002 | 0.003 | 0.003 | 0.000 | NS | 0.000 |
| *Elusimicrobia* | Lineage_IIa | 0.010 | 0.075 | 0.042 | 0.000 | 0.000 | 0.000 | 0.045 | 0.020 | 0.033 | 0.000 | NS | 0.000 |
| *Armatimonadetes* | *Fimbriimonadia* | 0.074 | 0.037 | 0.055 | 0.008 | 0.000 | 0.004 | 0.006 | 0.003 | 0.005 | 0.000 | NS | 0.000 |
| *Cyanobacteria* | *Melainabacteria* | 0.041 | 0.034 | 0.038 | 0.000 | 0.000 | 0.000 | 0.024 | 0.024 | 0.024 | 0.000 | NS | 0.000 |
| *Patescibacteria* | *Gracilibacteria* | 0.081 | 0.019 | 0.050 | 0.006 | 0.000 | 0.003 | 0.007 | 0.000 | 0.004 | 0.002 | NS | 0.002 |
| *Chloroflexi* | *Dehalococcoidia* | 0.020 | 0.028 | 0.024 | 0.000 | 0.000 | 0.000 | 0.047 | 0.011 | 0.029 | 0.000 | NS | 0.000 |
| *Planctomycetes* | vadinHA49 | 0.028 | 0.024 | 0.026 | 0.007 | 0.000 | 0.004 | 0.029 | 0.011 | 0.020 | 0.000 | NS | 0.000 |
| *Patescibacteria* | ABY1 | 0.016 | 0.046 | 0.031 | 0.000 | 0.000 | 0.000 | 0.021 | 0.002 | 0.012 | 0.000 | NS | 0.000 |
| *Chloroflexi* | AD3 | 0.001 | 0.002 | 0.001 | 0.006 | 0.000 | 0.003 | 0.075 | 0.000 | 0.037 | 0.000 | NS | 0.000 |
| WS2 | WS2_cl | 0.040 | 0.016 | 0.028 | 0.015 | 0.000 | 0.008 | 0.005 | 0.004 | 0.005 | 0.000 | NS | 0.000 |
| *Patescibacteria* | *Berkelbacteria* | 0.004 | 0.007 | 0.005 | 0.061 | 0.000 | 0.030 | 0.001 | 0.003 | 0.002 | 0.000 | NS | 0.000 |
| *Chloroflexi* | JG30-KF-CM66 | 0.005 | 0.008 | 0.007 | 0.000 | 0.000 | 0.000 | 0.028 | 0.030 | 0.029 | 0.000 | NS | 0.000 |
| *Acidobacteria* | Subgroup_22 | 0.002 | 0.049 | 0.026 | 0.008 | 0.000 | 0.004 | 0.001 | 0.003 | 0.002 | 0.000 | NS | 0.000 |
| *Cyanobacteria* | *Sericytochromatia* | 0.005 | 0.015 | 0.010 | 0.000 | 0.000 | 0.000 | 0.005 | 0.034 | 0.019 | 0.000 | NS | 0.000 |
| *Firmicutes* | *Erysipelotrichia* | 0.001 | 0.002 | 0.002 | 0.000 | 0.000 | 0.000 | 0.055 | 0.000 | 0.027 | 0.000 | NS | 0.000 |
| *Actinobacteria* | *Rubrobacteria* | 0.008 | 0.005 | 0.007 | 0.000 | 0.000 | 0.000 | 0.022 | 0.020 | 0.021 | 0.002 | NS | 0.002 |
| *Patescibacteria* | WWE3 | 0.001 | 0.045 | 0.023 | 0.000 | 0.000 | 0.000 | 0.002 | 0.000 | 0.001 | 0.000 | NS | 0.000 |
| *Armatimonadetes* | *Armatimonadia* | 0.004 | 0.000 | 0.002 | 0.000 | 0.000 | 0.000 | 0.017 | 0.022 | 0.019 | 0.000 | NS | 0.000 |
| *Chloroflexi* | SHA-26 | 0.002 | 0.019 | 0.011 | 0.000 | 0.000 | 0.000 | 0.011 | 0.011 | 0.011 | 0.000 | NS | 0.000 |
| *Fibrobacteres* | *Fibrobacteria* | 0.007 | 0.006 | 0.006 | 0.019 | 0.000 | 0.009 | 0.011 | 0.000 | 0.005 | 0.000 | NS | 0.000 |
| *Gemmatimonadetes* | BD2- | 0.029 | 0.008 | 0.019 | 0.000 | 0.000 | 0.000 | 0.003 | 0.000 | 0.002 | 0.000 | NS | 0.000 |
| *Acidobacteria* | Subgroup_25 | 0.002 | 0.026 | 0.014 | 0.000 | 0.000 | 0.000 | 0.012 | 0.000 | 0.006 | 0.000 | NS | 0.000 |
| *Gemmatimonadetes* | *Longimicrobia* | 0.032 | 0.001 | 0.016 | 0.006 | 0.000 | 0.003 | 0.000 | 0.000 | 0.000 | 0.000 | NS | 0.000 |
| *Elusimicrobia* | *Elusimicrobia* | 0.003 | 0.024 | 0.014 | 0.000 | 0.000 | 0.000 | 0.001 | 0.002 | 0.002 | 0.000 | NS | 0.000 |
| *Latescibacteria* | *Latescibacteria_cl* | 0.001 | 0.020 | 0.010 | 0.000 | 0.000 | 0.000 | 0.006 | 0.003 | 0.005 | 0.000 | NS | 0.000 |
| *Fusobacteria* | *Fusobacteriia* | 0.000 | 0.000 | 0.000 | 0.000 | 0.022 | 0.011 | 0.000 | 0.000 | 0.000 | 0.000 | NS | 0.000 |
| *Omnitrophicaeota* | *Omnitrophicaeota_cl* | 0.010 | 0.006 | 0.008 | 0.000 | 0.000 | 0.000 | 0.005 | 0.000 | 0.003 | 0.000 | NS | 0.000 |
| *Elusimicrobia* | Lineage_I Ib | 0.001 | 0.013 | 0.007 | 0.000 | 0.000 | 0.000 | 0.002 | 0.006 | 0.004 | 0.000 | NS | 0.000 |
| *Firmicutes* | *Erysipelotrichia* | 0.000 | 0.000 | 0.000 | 0.000 | 0.000 | 0.000 | 0.013 | 0.008 | 0.011 | 0.000 | NS | 0.000 |
| *Planctomycetes* | Pla4_lineage | 0.003 | 0.012 | 0.007 | 0.000 | 0.000 | 0.000 | 0.001 | 0.004 | 0.003 | 0.000 | NS | 0.000 |
| *Hydrogenedentes* | *Hydrogenedentia* | 0.003 | 0.005 | 0.004 | 0.000 | 0.000 | 0.000 | 0.001 | 0.004 | 0.003 | 0.000 | NS | 0.000 |
| *Omnitrophicaeota* | *Omnitrophia* | 0.001 | 0.004 | 0.002 | 0.000 | 0.000 | 0.000 | 0.008 | 0.000 | 0.004 | 0.000 | NS | 0.000 |
| *Acidobacteria* | Subgroup_15 | 0.007 | 0.000 | 0.004 | 0.000 | 0.000 | 0.000 | 0.001 | 0.004 | 0.003 | 0.000 | NS | 0.000 |
| *Planctomycetes* | BD7-11 | 0.001 | 0.001 | 0.001 | 0.000 | 0.000 | 0.000 | 0.004 | 0.006 | 0.005 | 0.000 | NS | 0.000 |
| *Bacteroidetes* | *Rhodothermia* | 0.000 | 0.000 | 0.000 | 0.000 | 0.000 | 0.000 | 0.000 | 0.000 | 0.000 | 0.011 | NS | 0.011 |
| *Acidobacteria* | AT-s3-28 | 0.001 | 0.003 | 0.002 | 0.000 | 0.000 | 0.000 | 0.004 | 0.001 | 0.003 | 0.000 | NS | 0.000 |
| *Patescibacteria* | *Kazania* | 0.009 | 0.000 | 0.005 | 0.000 | 0.000 | 0.000 | 0.000 | 0.000 | 0.000 | 0.000 | NS | 0.000 |
| *Entotheonellaeota* | *Entotheonellia* | 0.002 | 0.005 | 0.004 | 0.000 | 0.000 | 0.000 | 0.000 | 0.000 | 0.000 | 0.000 | NS | 0.000 |
| FCPU426 | FCPU426_cl | 0.002 | 0.002 | 0.002 | 0.000 | 0.000 | 0.000 | 0.003 | 0.000 | 0.002 | 0.000 | NS | 0.000 |
| *Actinobacteria* | FFCH16263 | 0.002 | 0.004 | 0.003 | 0.000 | 0.000 | 0.000 | 0.000 | 0.000 | 0.000 | 0.000 | NS | 0.000 |

NS: No sample

**Table S7:**
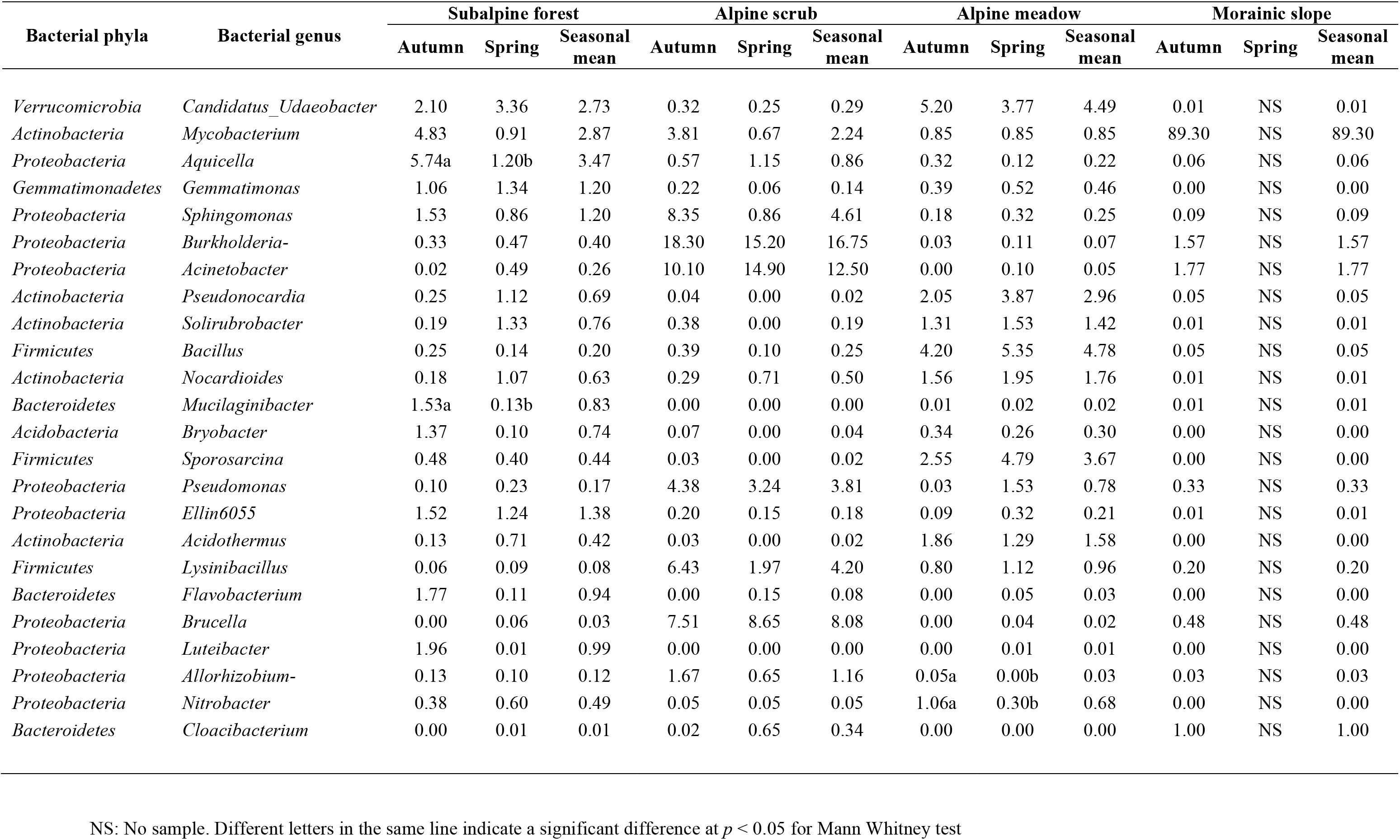
Relative abundance of abundant bacterial genus (≥ 1% relative abundances) found in the four habitats in autumn and spring.

**Table S8:**
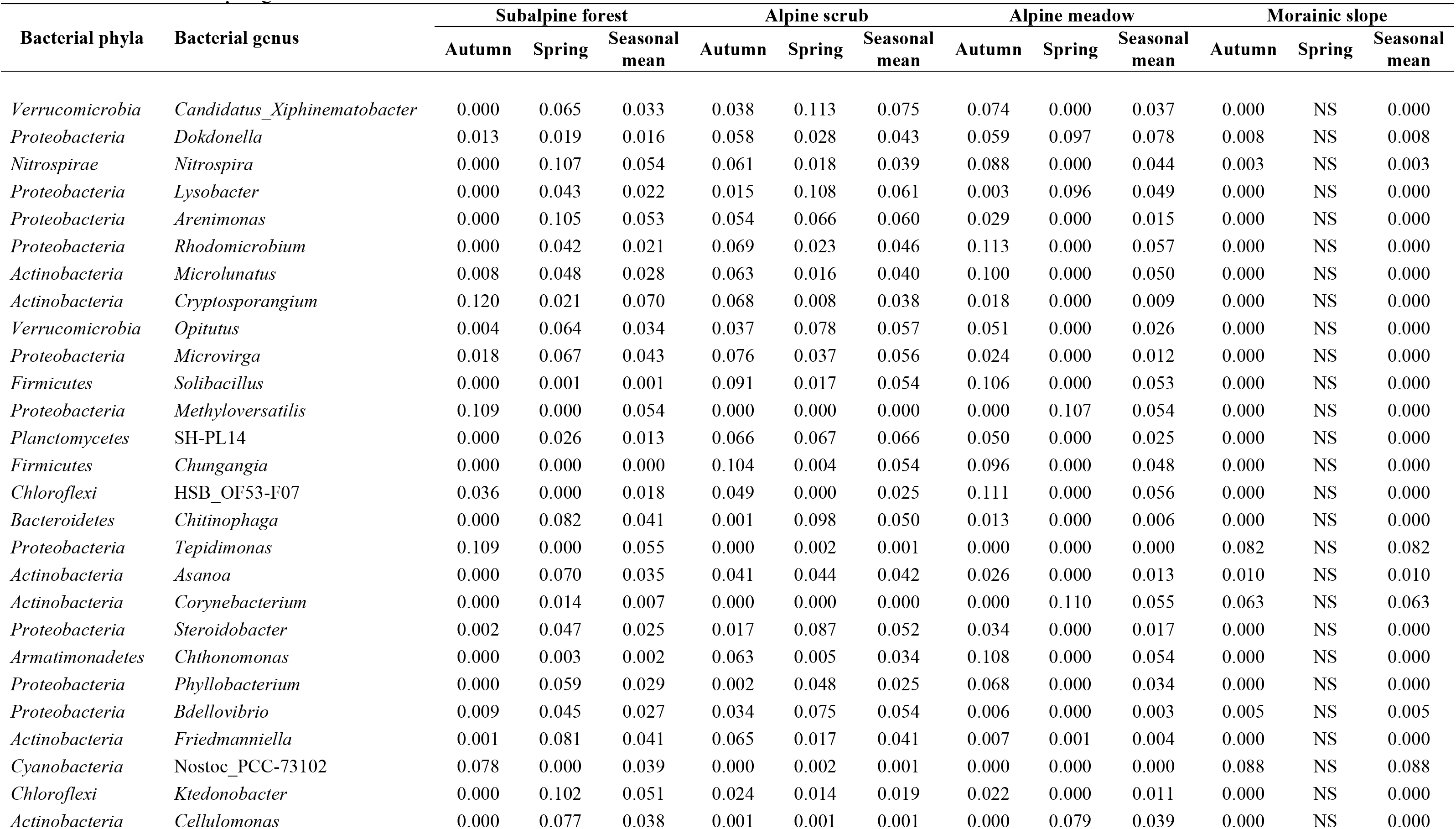
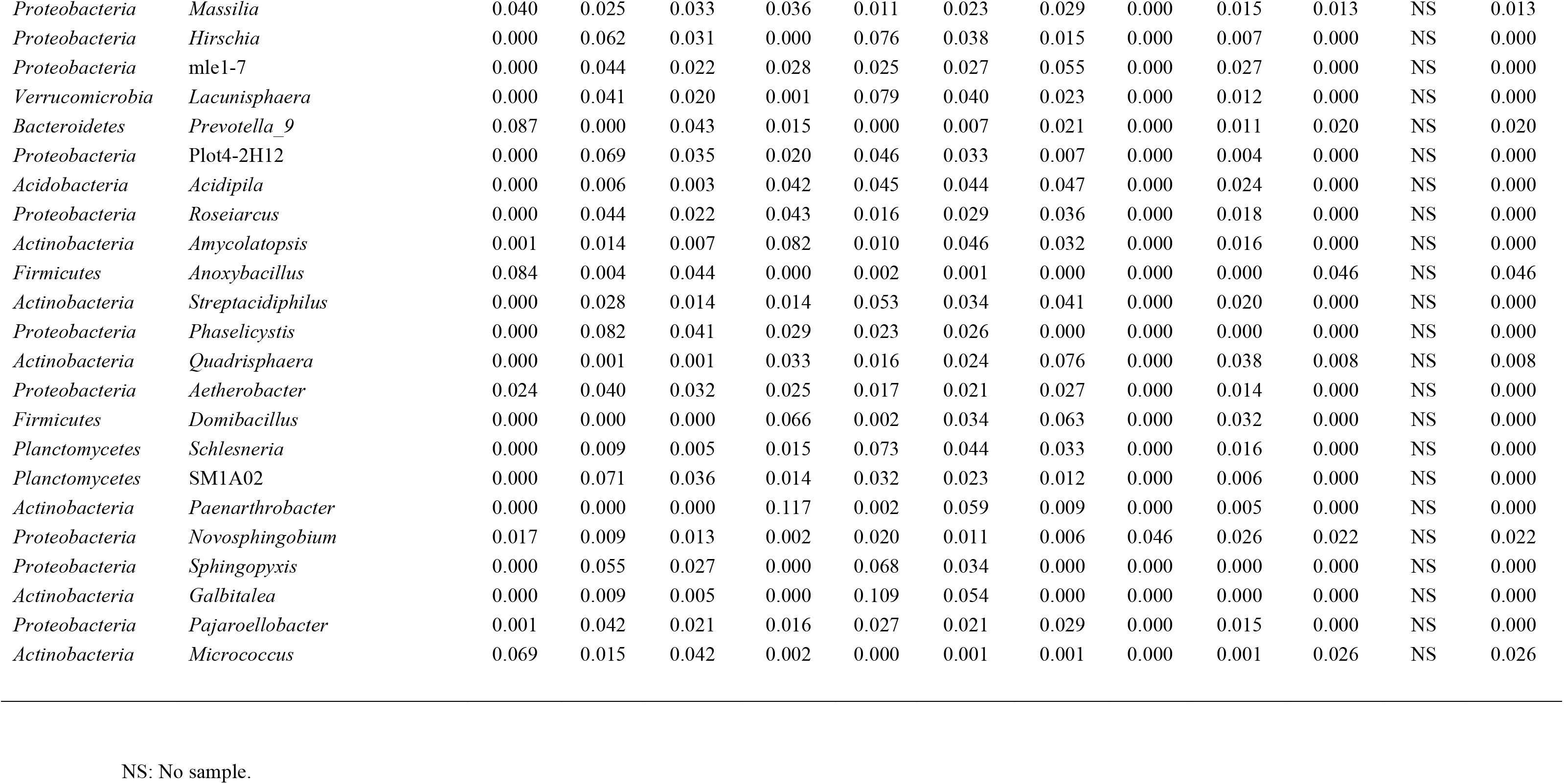
Relative abundance of 50 most abundant rare bacterial genus (≤ 0.1% relative abundances) found in the four habitats in autumn and spring.

**Table S9:** Results of MRPP analysis for bacterial community composition comparison between four habitats along elevation.

| Bacterial community | Overall |  | Habitats<sup>b</sup><br>Comparison pairs | Pairwise |  |
| --- | --- | --- | --- | --- | --- |
|  | A value<sup>a</sup> | *p*-value |  | A value<sup>a</sup> | *p*-value |
| All genera | **0.3114** | **0.0001** | SF vs AS | **0.184** | **0.0001** |
|  |  |  | SF vs AM | **0.1365** | **0.0002** |
|  |  |  | SF vs MS | **0.3039** | **0.0004** |
|  |  |  | AS vs AM | **0.2568** | **0.0001** |
|  |  |  | AS vs MS | **0.176** | **0.0002** |
|  |  |  | AM vs MS | **0.4316** | **0.0008** |
| Abundant genera | **0.4693** | **0.0001** | SF vs AS | **0.2956** | **0.0001** |
|  |  |  | SF vs AM | **0.2229** | **0.0001** |
|  |  |  | SF vs MS | **0.3698** | **0.0005** |
|  |  |  | AS vs AM | **0.4377** | **0.0001** |
|  |  |  | AS vs MS | **0.4378** | **0.0004** |
|  |  |  | AM vs MS | **0.6029** | **0.0007** |
| Rare genera | **0.1501** | **0.0001** | SF vs AS | **0.1214** | **0.0009** |
|  |  |  | SF vs AM | **0.1436** | **0.0025** |
|  |  |  | SF vs MS | **0.198** | **0.001** |
|  |  |  | AS vs AM | **0.1645** | **0.0057** |
|  |  |  | AS vs MS | 0.01026 | 0.1555 |
|  |  |  | AM vs MS | **0.2728** | **0.0065** |

<sup>a</sup>A-value: Chance corrected within-group agreement
<sup>b</sup>Habitats: SF, Subalpine forest; AS, Alpine scrub; AM, Alpine meadow; MS, Morainic slope
A values and *p*-values in bold indicate significant differences ( $p < 0.05$ )

**Table S10:** Average dissimilarity in the bacterial communities at (a) phyla level (b) class level (c) genus level between the four habitats as obtained from SIMPER analysis.

| **Average dissimilarity** | **Alpine scrub** | **Alpine meadow** | **Morainic slope** |
| --- | --- | --- | --- |
| **Subalpine forest** | 32.6% | 29.6% | 44.0% |
| **Alpine scrub** |  | 37.0% | 40.7% |
| **Alpine meadow** |  |  | 38.6% |

(b)
| **Average dissimilarity** | **Alpine scrub** | **Alpine meadow** | **Morainic slope** |
| --- | --- | --- | --- |
| **Subalpine forest** | 40.0% | 34.0% | 48.0% |
| **Alpine scrub** |  | 42.0% | 45.0% |
| **Alpine meadow** |  |  | 46.0% |

(c)
| **Average dissimilarity** | **Alpine scrub** | **Alpine meadow** | **Morainic slope** |
| --- | --- | --- | --- |
| **Subalpine forest** | 69.0% | 49.0% | 67.0% |
| **Alpine scrub** |  | 68.0% | 69.0% |
| **Alpine meadow** |  |  | 64.0% |

**Table S11:** Results of MRPP analysis for bacterial community composition comparison between autumn and spring.

| Bacterial community | Habitats<sup>b</sup> | Pairwise |  |
| --- | --- | --- | --- |
|  |  | A value<sup>a</sup> | *p*-value |
| All genera | SF | 0.003429 | 0.453 |
|  | AS | 0.02959 | 0.2249 |
|  | AM | -0.00483 | 0.5146 |
| Abundant genera | SF | 0.01773 | 0.35 |
|  | AS | 0.03098 | 0.2679 |
|  | AM | -0.02974 | 0.5701 |
| Rare genera | SF | -0.009749 | 0.6082 |
|  | AS | -0.005276 | 0.6583 |
|  | AM | 0.01057 | 0.2573 |

<sup>a</sup>A-value: Chance corrected within-group agreement
<sup>b</sup>Habitats: SF, Subalpine forest; AS, Alpine scrub; AM, Alpine meadow

**Table S12:** Mantel test results for the correlation between bacterial community composition and environmental variables along elevation gradient.

|  | Elevation |  | MT<sub>30</sub> |  | SMC |  | pH |  | SOC |  | TN |  | C/N |  |
| --- | --- | --- | --- | --- | --- | --- | --- | --- | --- | --- | --- | --- | --- | --- |
|  | r | p-value | r | p-value | r | p-value | r | p-value | r | p-value | r | p-value | r | p-value |
| **Bacterial community** |  |  |  |  |  |  |  |  |  |  |  |  |  |  |
| **All phyla** | **0.4043** | **0.0001** | **0.3587** | **0.0001** | 0.0519 | 0.1787 | **0.2454** | **0.0013** | **0.2697** | **0.0004** | **0.3273** | **0.0001** | **0.3249** | **0.0001** |
| **Abundant phyla** | **0.4056** | **0.0001** | **0.3592** | **0.0001** | 0.0509 | 0.1910 | **0.2448** | **0.0019** | **0.2689** | **0.0004** | **0.3287** | **0.0001** | **0.3226** | **0.0001** |
| **Rare Phyla** | 0.0606 | 0.1950 | -0.0518 | 0.6895 | 0.1091 | 0.1517 | **0.1928** | **0.0199** | 0.1702 | 0.0584 | 0.1550 | 0.0794 | 0.1172 | 0.1481 |
| **All Class** | **0.3587** | **0.0001** | **0.3520** | **0.0001** | 0.0534 | 0.1764 | **0.2642** | **0.0013** | **0.3456** | **0.0001** | **0.3826** | **0.0001** | **0.3592** | **0.0001** |
| **Abundant Class** | **0.3593** | **0.0001** | **0.3505** | **0.0001** | 0.0492 | 0.1970 | **0.2658** | **0.0009** | **0.3437** | **0.0002** | **0.3832** | **0.0001** | **0.3602** | **0.0001** |
| **Rare class** | 0.1530 | 0.0630 | 0.1501 | 0.0686 | -0.0236 | 0.5482 | **0.3492** | **0.0028** | **0.4259** | **0.0001** | **0.3785** | **0.0001** | **0.3115** | **0.0048** |
| **All genera** | **0.2878** | **0.0013** | **0.2175** | **0.0079** | 0.0930 | 0.1003 | **0.2657** | **0.0029** | **0.3801** | **0.0003** | **0.4607** | **0.0001** | **0.2812** | **0.0015** |
| **Abundant genera** | **0.2775** | **0.0013** | **0.2225** | **0.0069** | 0.1038 | 0.0773 | **0.2781** | **0.0015** | **0.3501** | **0.0001** | **0.4170** | **0.0001** | **0.2532** | **0.0006** |
| **Rare genera** | 0.1115 | 0.1189 | 0.0986 | 0.1526 | -0.0484 | 0.6275 | **0.2404** | **0.0225** | **0.1767** | **0.0326** | **0.2931** | **0.0014** | **0.2219** | **0.0519** |

p-values in bold indicate significant correlation
MT<sub>30</sub>, mean temperature of 30 days period; SMC, soil moisture content; SOC, soil organic carbon, TN, total nitrogen; C/N, carbon nitrogen ratio

**Table S13:** Results of Univariate regression representing relationship of relative abundances of abundant bacterial class with environmental variables.

| Bacterial phyla | MT<sub>30</sub> |  |  | TN |  |  | SOC |  |  |
| --- | --- | --- | --- | --- | --- | --- | --- | --- | --- |
|  | R-squared | Estimate<sup>a</sup> | p-value<sup>b</sup> | R-squared | Estimate | p-value | R-squared | Estimate | p-value |
| *Proteobacteria* | 0.275 | 0.045 | 0.0008*** | - | - | - | - | - | - |
| *Actinobacteria* | 0.32 | -0.051 | 0.00025*** | - | - | - | - | - | - |
| *Firmicutes* | - | - | - | 0.2213 | 0.048 | 0.00293** | 0.1078 | 0.0026 | 0.05* |
| *Patescibacteria* | 0.3379 | 0.023812 | 0.000186*** | - | - | - | - | - | - |
| *Bacteroidetes* | 0.1648 | 0.016525 | 0.00995** | - | - | - | - | - | - |
| *Verrucomicrobia* | - | - | - | 0.4519 | 0.03091 | 0.0000133** | 0.4402 | 0.002499 | 0.000019** |
| *Acidobacteria* | - | - | - | 0.244 | 0.0022 | 0.00299** | 0.2011 | 0.025 | 0.00781** |
| *Planctomycetes* | - | - | - | 0.52 | 0.03803 | 0.00000146* | 0.411 | 0.0027 | 0.00044*** |
| *Chlamydiae* | 0.2172 | 0.0133 | 0.00548** | - | - | - | - | - | - |
| *Gemmatimonadetes* | - | - | - | - | - | - | - | - | - |
| *Dependentiae* | - | - | - | - | - | - | - | - | - |
| *Chloroflexi* | - | - | - | 0.2601 | 0.0016 | 0.00206** | 0.278 | 0.02 | 0.00136** |

MT<sub>30</sub>, mean temperature of 30 days period; TN, total nitrogen; SOC, soil organic carbon
<sup>a</sup>Estimate: $\beta$ coefficient
<sup>b</sup>p-value (significance value): \*\*\*, P < 0.001; \*\*, P < 0.01; \*, P < 0.05

**Table S14:** Results of Univariate regression representing relationship of relative abundances of abundant bacterial class with environmental variables.

| Bacterial phyla | Bacterial class | MT<sub>30</sub> |  |  | TN |  |  | SOC |  |  |
| --- | --- | --- | --- | --- | --- | --- | --- | --- | --- | --- |
|  |  | R-squared | Estimate<sup>a</sup> | p-value<sup>b</sup> | R-squared | Estimate | p-value | R-squared | Estimate | p-value |
| *Proteobacteria* | *Gammaproteobacteria* | 0.168 | 0.03 | 0.016* | - | - | - | - | - | - |
| *Proteobacteria* | *Alphaproteobacteria* | 0.3451 | 0.031119 | 0.00026*** | - | - | - | - | - | - |
| *Proteobacteria* | *Deltaproteobacteria* | - | - | - | 0.2912 | 0.012142 | 0.000989*** | 0.3009 | 0.0010116 | 0.000782*** |
| *Actinobacteria* | *Actinobacteria* | 0.3563 | -0.0534 | 0.000194*** | - | - | - | - | - | - |
| *Actinobacteria* | *Thermoleophilia* | - | - | - | 0.367 | 0.036553 | 0.000147*** | 0.3209 | 0.0028011 | 0.000479*** |
| *Actinobacteria* | *Acidimicrobiia* | - | - | - | 0.3832 | 0.016836 | 0.000095*** | 0.4171 | 0.0014396 | 0.0000371*** |
| *Firmicutes* | *Bacilli* | - | - | - | 0.2582 | 0.05071 | 0.00216** | 0.1159 | 0.002785 | 0.0488* |
| *Patescibacteria* | *Saccharimonadia* | 0.3616 | 0.023621 | 0.000169*** | - | - | - | - | - | - |
| *Bacteroidetes* | *Bacteroidia* | 0.1931 | 0.016525 | 0.00932** | - | - | - | - | - | - |
| *Verrucomicrobia* | *Verrucomicrobiae* | - | - | - | 0.4466 | 0.03175 | 0.0000156** | 0.4333 | 0.0025631 | 0.0000232*** |
| *Acidobacteria* | *Subgroup_6* | - | - | - | 0.1356 | 0.0164 | 0.0321* | 0.162 | 0.0014691 | 0.0183* |
| *Acidobacteria* | *Acidobacteriia* | - | - | - | 0.2063 | 0.016395 | 0.00697** | 0.2562 | 0.001497 | 0.00226** |
| *Acidobacteria* | *Blastocatellia_Subgroup_4* | - | - | - | 0.3729 | 0.015414 | 0.000125*** | 0.3475 | 0.0012194 | 0.000244*** |
| *Planctomycetes* | *Planctomycetacia* | - | - | - | 0.4922 | 0.032076 | 0.00000379* | 0.3953 | 0.0023558 | 0.0000683*** |
| *Planctomycetes* | *Phycisphaerae* | - | - | - | 0.4868 | 0.020527 | 0.00000451* | 0.358 | 0.0014427 | 0.000186*** |
| *Chlamydiae* | *Chlamydiae* | 0.2273 | 0.013956 | 0.00436** | - | - | - | - | - | - |
| *Gemmatimonadetes* | *Gemmatimonadetes* | - | - | - | - | - | - | - | - | - |
| *Dependentiae* | *Babeliae* | - | - | - | - | - | - | - | - | - |
| *Chloroflexi* | *KD4-96* | - | - | - | 0.2055 | 0.012376 | 0.00709** | 0.2225 | 1.06E-03 | 0.00486** |

MT<sub>30</sub>, mean temperature of 30 days period; TN, total nitrogen; SOC, soil organic carbon
<sup>a</sup>Estimate: $\beta$ coefficient
<sup>b</sup>p-value (significance value): \*\*\*, P < 0.001; \*\*, P < 0.01; \*, P < 0.05

## Notes

### Competing Interest Statement

The authors have declared no competing interest.

